# System Analysis of the Sequencing Quality of Human Whole Exome Samples on Bgi Ngs Platform

**DOI:** 10.1101/2021.02.02.429394

**Authors:** Vera Belova, Anna Pavlova, Robert Afasizhev, Viktoria Moskalenko, Margarita Korzhanova, Valery Cheranev, Andrey Krivoy, Boris Nikashin, Irina Bulusheva, Denis Rebrikov, Dmitriy Korostin

**Affiliations:** Center for Precision Genome Editing and Genetic Technologies for Biomedicine, Pirogov Medical University, 117997 Moscow, Russia

## Abstract

Human whole exome sequencing (WES) is now the standard for most medical genetics applications worldwide. The leaders are manufacturers of enrichment kits that base their protocols on a hybridization approach using cRNA or cDNA biotinylated samples specific to regions of interest in the genome. Recently, platforms from the Chinese company MGI Tech have been successfully promoted in the markets of many countries in Europe and Asia. There is no longer any question about their reliability and the quality of the data obtained. However, very few task-specific kits for WES, in particular, are presented for these sequencers. We have developed our solution for library pre-capture pooling and exome enrichment using Agilent probes. In this work, we demonstrate on a set of standard benchmark samples from the Platinum Genome Collection that our protocol, called “RSMU_exome”, is superior to the kit from MGI Tech in qualitative and quantitative terms. It allows detecting more SNVs and CNVs with superior sensitivity and specificity values, generates fewer PCR duplicates, allows more samples to be pooled in a single enrichment, and requires less raw data to produce results comparable to the MGI Tech solution. Also, our protocol is significantly cheaper than the kit from the Chinese manufacturer.

## Introduction

Combining diagnostic and research capabilities, WES is today the most important method for studying hereditary pathology. At the current level of science and technology development for solving research and clinical diagnostic problems, whole-exome sequencing has several advantages over whole-genome sequencing. WES allows us to focus on the study of the most clinically valuable part of the human genome - protein-coding sequences. It is known that approximately 85% of the genetic variants responsible for the development of human inherited diseases are found in exons and intronic splicing sites [1].

Also, the economic factor continues to play an important role in the introduction of genomic technologies into clinical practice. Unlike whole-genome sequencing, in which ~3 billion bp of the human genome must be read, whole exome sequencing involves capturing and targeting the coding and adjacent regions, which corresponds to 1-2% of the human genome. On average over the past decade, the WES for a patient has been 4-5 times cheaper than the WGS test [2]. At the same time, WGS wins only 1-2% over WES in the efficiency of diagnosis, because only a small number of pathogenic variants registered in ClinVar are not detected by known exome kits [3, 4].

Most exome enrichment solutions have been developed for Illumina sequencers, the best known being SureSelect (Agilent), SeqCap EZ (Roche NimbleGen), and TruSeq Capture (Illumina). The essence of the method is the hybridization of biotinylated DNA or RNA probes with complementary exome fragments of DNA libraries. Enrichment kits differ in the size of the target region, the length, type and density of the probes, as well as the number of samples enriched in one reaction [5]. Since each manufacturer tries to improve its method from year to year, new comparisons of the kits with each other are periodically published [6-10]. First of all, the comparison is made according to the following parameters of interest to researchers: target enrichment efficiency, uniformity of coverage, sequencing complexity and ability to call true SNVs. Currently, the major NGS platform in the world is Illumina sequencer producing almost 90% of sequencing data [11]. At the end of 2017, Chinese company MGI Tech presented MGISEQ-2000 sequencing platform promoting it as a device for large and medium scale genome sequencing. MGISEQ is specific in harnessing cPAS sequencing technology and using nanoballs (DNB) generated from circular molecules of DNA library by rolling circle replication [12]. There are a few research articles published about the performance of MGISEQ-2000 in recent 2 years but most of them state that this new sequencer is not inferior in sequencing quality to Illumina machines [13-16].

There are 2 commercial exome enrichment kits for the MGISEQ-2000 (now DNBSEQ-G400), MGI Tech’s own MGIEasy Exome Capture V4 Probe Set and MGIEasy Exome Capture V5 Probe Set, which differ only in different probe versions, but not in the enrichment protocol. We tested the first MGIEasy Exome Capture V4 Probe Set released in 2019 using the benchmark sample NA12891 from the Platinum Genomes project [17].

In the absence of a large selection of kits and protocols for the brand-new MGISEQ-2000 sequencer we also present our improved hybridization and capture method for WES. We showed performance and compatibility of our custom “RSMU_exome” protocol with MGIEasy v4 probes and Agilent SureSelect All Exon v6 probes (hereinafter MGIEasy v4 and Agilent v6) (**Error! Reference source not found.Error! Reference source not found.Error! Reference source not found.**). We made 2 pools A and B of 24 independently prepared human gDNA libraries - 10-12 libraries in each pool, both of which were enriched in 3 different ways - by our custom “RSMU_exome” protocol with Agilent v6 probes and separately with MGIEasy v4 probes and by the standard MGIEasy Exome Capture V4 Probe Set protocol for comparison.

**Figure 1.**
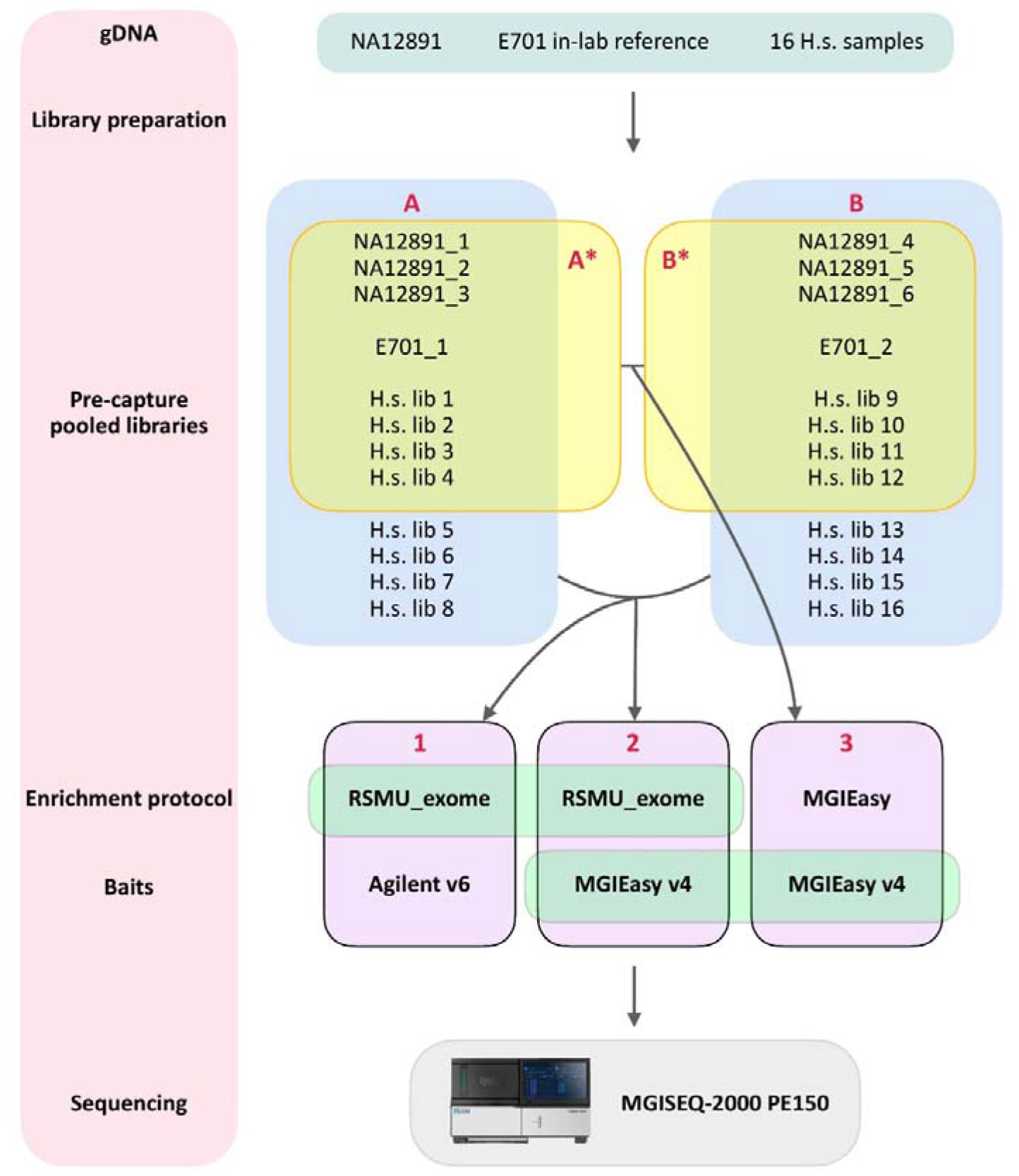
Experiment scheme. We made two pools, A and B, of 24 independently prepared human gDNA libraries - in each pool 8 or 12 libraries, of which 3 DNA libraries belonged to sample NA12891 (a total of 6 independent libraries for NA12891) and 1 library to the in-lab reference sample E701. Both pools were independently enriched in 3 different ways - using our own “RSMU_exome” protocol with Agilent v6 probes, then separately with MGIEasy v4 probes and using the standard MGIEasy Exome Capture V4 Probe Set for comparison. Thus, in total, we enriched 6 pools (64 libraries in total) and obtained a data set consisting of 64 pairs of fastq files.

## Materials and Methods (Belova, Pavlova, Afasizhev)

### Ethics Statement

This study was conducted according to the principles expressed in the Declaration of Helsinki. Appropriate institutional review board approval for this study was obtained from the Ethics Committee at Pirogov Medical University. All patients provided written informed consent for the collection of samples, subsequent analysis, and publication thereof.

### Sample collection

For the study, we used a reference DNA sample NA12891. We also isolated DNA from blood samples from 16 patients, and DNA from our in-laboratory patient reference blood sample.

### DNA extraction

DNA isolation was performed using the DNeasy Blood and Tissue Kit according to the manufacturer’s protocol (Qiagen). Extracted DNA was then quantified with a Qubit dsDNA BR Assay system (Life Technologies) and the quality of DNA was assessed on 1% agarose gel.

### Library preparation

Then we prepared 1 library for each of 16 DNA patients’ samples, 2 independent libraries for our in-lab reference sample E701 and 6 independent libraries for NA12891 sample. For each library, 400 ng of input genomic DNA was sheared to ~300 bp with Covaris E220 System according to recommended manufacturer’s procedure. Then size-selection of DNA fragments was performed by SPRI AMPure XP beads (Beckman Coulter) to achieve a target peak of 240 to 290 bp (first cut ratio x0,8, second cut ratio x0,2).

Further steps of library preparation were proceeded using MGIEasy Universal DNA Library Prep Set (MGITech), except for PCR step as described below. For the further pooling of individual libraries, balanced combinations of barcoded adapters from the MGIEasy DNA Adapters-96 (Plate) Kit were selected for the libraries using the BC-store software developed in our laboratory [18] according to strict criteria. Thus, each library was ligated to adapter containing one selected barcode. To amplify libraries, we used KAPA Hi-Fi polymerase (KAPA Biosystems) instead of MGI polymerase according to the manufacturer’s recommendations with the following PCR program: a 3 min activation step at 95°C, followed by 10-12 cycles of 20 s at 98°C, 15 s at 60°C, 30 s at 72°C, with a final extension of 10 min at 72°C. The quality control of the obtained DNA libraries was carried out with the gel electrophoresis and with the High Sensitivity DNA assay on the 2100 Bioanalyzer System (Agilent Technologies). Library peak size was in the range of 300 to 400 nucleotides. Libraries concentrations then were quantified by fluorometry with a Qubit dsDNA HS Assay system (Life Technologies). The total amount of each DNA library had to be> = 1100 ng to be sufficient for 3 different enrichment procedures.

### Pre-capture sample pooling

We designed 2 pools of libraries “A” and “B” (**Error! Reference source not found.**), consisting of 12 different libraries each, 3 of which belonged to the NA12891 reference sample, 1 to the E701 our in-lab reference sample and 8 to the patient collection DNA samples.

Both pools “A” and “B” went into three separate paths:

1. Exome enrichment with MGIEasy protocol with MGIEasy Exome Capture V4 Probe;
2. Exome enrichment with “RSMU_exome” protocol with MGIEasy Exome Capture V4 Probe;
3. Exome enrichment with “RSMU_exome” protocol with Agilent SureSelect All Exon v6 probe.

For the MGIEasy enrichment protocol (path 1), the pools were reduced to 8 libraries per pool according to the manufacturer’s recommendations [19]. For 8-plex hybridization, 250 ng of each library is required and the total amount of DNA in the pool was 2 μg.

For the “RSMU_exome” enrichment protocol (S1 File), we combined 400 ng input of each of the 12 libraries, so the total amount of DNA in the pool was 4.8 μg. The library pools were then completely dried on a speedVac concentrator (ThermoFisher) at 50 °C.

### Enrichment methods

#### MGIEasy Exome Capture

Pools A1 and B1 were hybridized and captured with MGIEasy Exome Capture V4 Probe following the manufacturer’s protocols. Hybridization was performed at 65°C for 24h then pools of libraries were captured using streptavidin-conjugated magnetic beads MyOne T1 Dynabeads on a room temperature with subsequent series of washes. Then post-capture amplification was performed with 13 cycles PCR.

### “RSMU_exome” protocol

#### Hybridization

Dried pools A2, B2 and A3, B3 were combined with 11 μL Cot-1 DNA (1 μg/μL, ThermoFisher) and 2 adaptor-blocking oligonucleotides with LNA modifications - 500 pmol each (Table 1). Samples were transferred to PCR tubes and denatured at 95°C for 5 min, followed by a second infinite hold at 65°C.

**Table 12.**
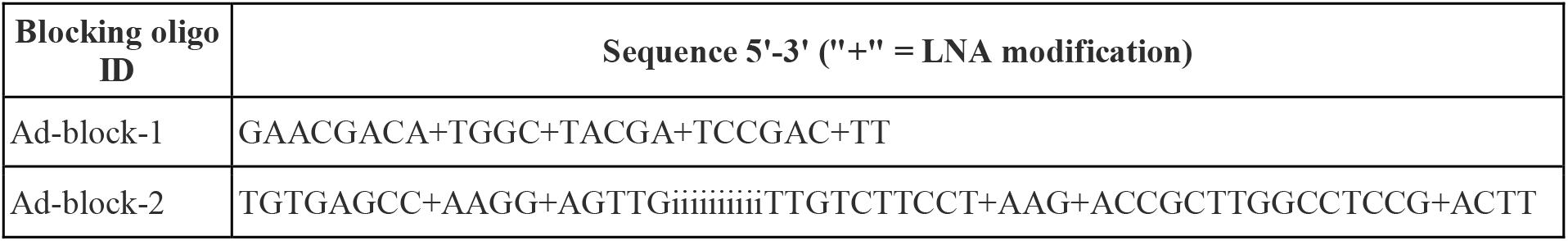
Blocking oligo sequences.

Then, to the samples we added 14 μL of hybridization buffer thoroughly vortexed and preheated at 65°C for 10 min to dissolve any precipitate (mix of components is listed in Table 3).

**Table 3.**
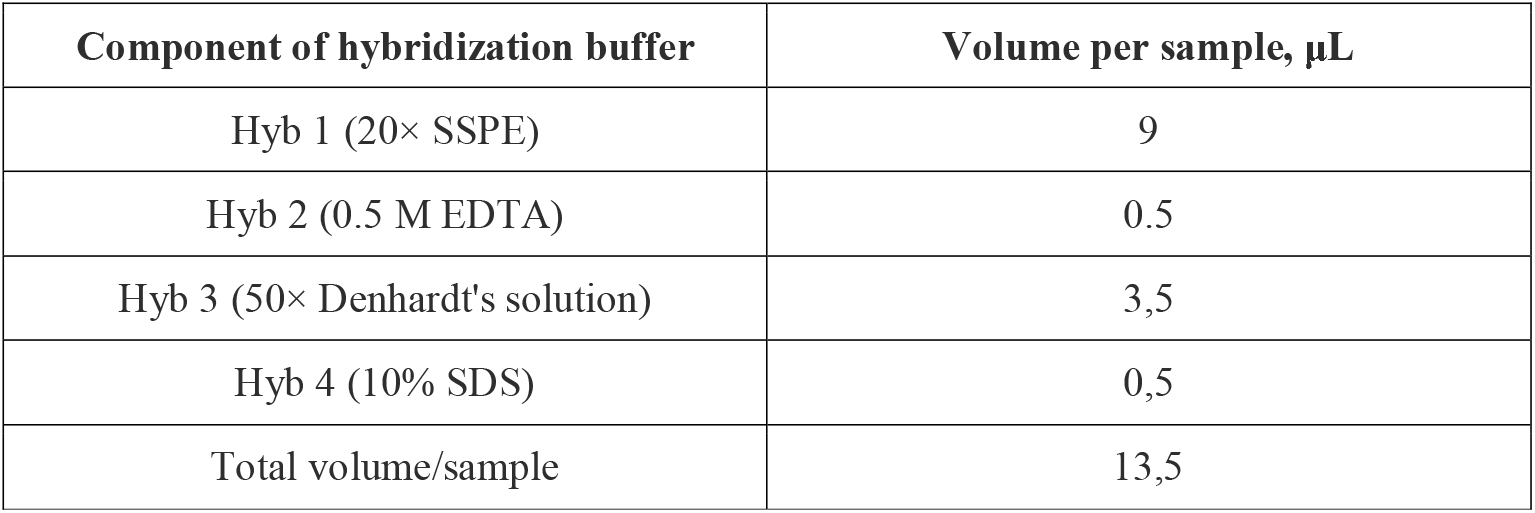
Components in hybridization buffer.

Finally mix of RNA baits (4 μL of Agilent v6 baits or 6 μL of MGI v4 baits) with 1 μL of SUPERase-In (20 U/μL, Invitrogen) blocker preheated at 65°C for 5 min were added to the samples right in the thermocycler. After slowly pipetting the samples, we added mineral oil to prevent its evaporation. We remain the hybridization mixture on the thermal cycler for 24 h at 65°C with a heated lid set at 105°C.

### Washing

Per 1 hybridization reaction, 3□μl C1 streptavidin Dynabeads (10 mg/mL, Invitrogen) were washed 3 times with 200 μL binding buffer (□1M NaCl, 10□mM Tris-HCl [pH 7.5], 1□mM EDTA) on a magnetic rack, then resuspended in 70 μL binding buffer in LoBind tubes (Eppendorf). After that, Dynabeads were incubated on a rotator for 15 min at room temperature with 3 μg/reaction of salmon sperm DNA. Prior to capture, we preheated Dynabeads at 65°C for 5 min.

Thereafter, the enriched pools were bound to streptavidin Dynabeads and left in a thermoshaker for 30 min at 65°C. Then, a series of 3 washes were carried out: we collected beads on a magnetic rack, removed supernatant, added 500 μL prewarmed (45 min at 65°C) wash buffer 0.02X SSC/0.01% SDS and transferred samples to the thermoshaker for 10 min at 65°C. After washes, samples were dried for 3-4 min on a magnetic stand and resuspended in 31 μL mQ in new PCR tubes.

Before amplification, we denatured enriched DNA libraries from Dynabeads by heating samples at 95°C 5 min and fast collecting supernatant on a magnetic rack to the new PCR tube.

### Amplification

The post capture PCR set-up was as follows: ½ volume of enriched pool e.g. 15mkl, 8 mkl 0.3 μM MGI primer mix (Table 4), 1,5 μL 10 mM dNTP Mix, 10μL of KAPA HiFi Fidelity buffer (5X) and 1 μL Kapa HiFi HotStart Polymerase (1 U/μL Kapa Biosystems) in a total volume of 50 μL. We used the following PCR program: 95°C for 3 min, 7 cycles × [98°C 20 s, 60°C 15 s, 72°C 30 s], 72°C 10 min. When MGI v4 baits were used for pools A2, B2, 10 cycles of PCR were completed for amplification.

**Table 45.**
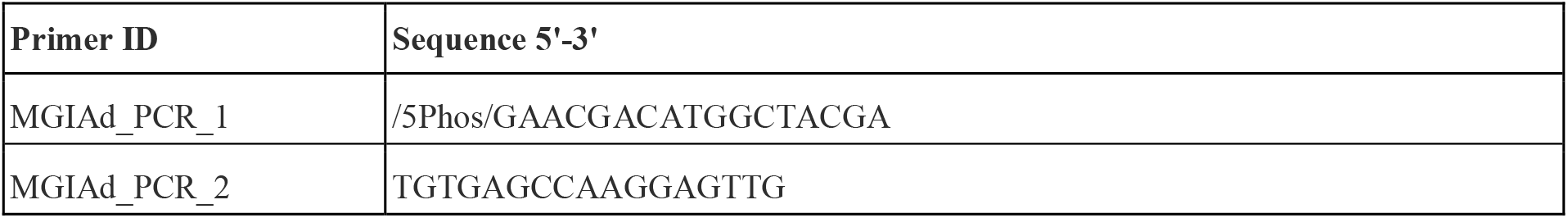
Primer sequences.

Immediately after PCR, the quality and size distribution of the enriched pools were checked by 2% gel electrophoresis, after which the entire volume of the PCR product was purified using 1x SPRI beads (Ampure) and eluted in 38 μL mQ water. Next, the concentration was measured using a Qubit dsDNA HS Assay system (Life Technologies).

### Sequencing

Finally, 6 pools of enriched DNA libraries were circularised to generate single stranded DNA circles. All samples were then processed for DNB generation and massive sequencing in Paired-end 2×150 bp mode according to MGI protocol. We loaded 1 pool per lane on the patterned flowcells in two different runs on MGISEQ-2000 platform.

### Bioinformatic pipeline

#### Data processing

The data processing scheme is shown in Figure 12**Error! Reference source not found.** The quality of the obtained fastq files was analyzed using FastQC v0.11.9 [20]. FastQC results were combined into a single report using multiqQC [21]. Based on quality, fastq files were trimmed using Trimmomatic v0.39 [22]. Reads were aligned to the indexed reference genome GRCh37 using bwa- mem [23]. SAM files were converted into BAM files and sorted using SAMtools v1.9 to check the percentage of aligned reads [24]. From the obtained bam files the quality metrics for exome enrichment and sequencing were collected using Picard v2.22.4 [25] and information on the number of duplicates was collected using Picard MarkDuplicates v2.22.4.

The resulting bam files were analyzed using two strategies: collecting WES enrichment quality metrics and calculating statistical enrichment parameters using NA12891 as the gold standard.

To correctly assess enrichment and sequencing quality, NA12891 samples were downsampled using Picard DownsampleSam v2.22.4 to 50 million reads. Then duplicates were removed from the downsampled samples with Picard MarkDuplicates v2.22.4, and Picard CollectHsMetrics v2.22.4 was used to collect quality statistics of the obtained data. To correctly compare the two enrichment reagent kits, we used the quality assessment of several bed files: MGIEasy v4, Agilent v6, the intersection of MGIEasy v4 and Agilent v6, and the set of all protein-coding regions by Ensembl.

The next step to assess the quality of the obtained data was to analyze the quality of SNV and CNV calling. Raw reads were aligned using the method described above. In the case of SNV and CNV analysis, first, we removed duplicates and then downsampled BAM files to 50 million reads. Variant calling was made for all samples from all pools using bcftools mpileup v1.9 and Intersection over Union (*IoU*) values were counted for all vcf files.

### Data availability

Fastq files for each library of NA12891 sample in all 6 pools were deposited in the NCBI open access sequence read archive (SRA) under BioProject ID PRJNA667840.

## Results (Belova, Pavlova, Afasizhev)

### Comparison of enrichment methods

We present the protocol for enrichment on different probes that we use in the lab for sequencing on the MGISEQ-2000. For comparison, we enriched the same library pools with two different protocols by ours and by MGI Tech, and showed that our protocol works well with both MGI Tech and Agilent probes.

Before enrichment, we suggest pooling 12 libraries (see S1 Table, S1 Figure for results for libraries in more detail), in the amount of 350-500 ng each; the MGI protocol suggests pooling a maximum of 8 libraries, at 250 ng each. The maximum number of DNA libraries per input with our “RSMU_exome” protocol is 5 μg, with the MGI protocol being 2 μg. Before capture, we block Dynabeads with Salmon sperm DNA since Dynabeads themselves are known to be capable of binding DNA: 1 mg of Dynabeads MyOne Streptavidin C1 typically binds ~20 μg ds-DNA and ~500 pmol ss-oligonucleotides [26]. This is how we prevent immobilization of DNA-RNA hybrids directly on the surface of Dynabeads. For the 3 washes, we use only one buffer and a constant temperature of 65°C. Before post-capture PCR, we denature samples DNA from Dynabeads, because in our experience (S2 Table) the efficiency of PCR with Dynabeads is ~25% lower than those without Dynabeads. We also use the high-processing Hi-Fi KAPA polymerase, which has proven to be the most efficient solution in library amplification experiments in our laboratory.

Despite the fact that in post-capture PCR we take only half of the reaction volume and use fewer cycles (7-10 cycles) than in the MGI Tech protocol (13 cycles), the yield of enriched pools of libraries in ng by our protocol is higher (Table 6), indicating a more efficient procedure of enrichment and amplification.

**Table 67.** Yield in ng of enriched pools of DNA libraries for different protocols.

| Pool ID | Protocol | Baits | input DNA lib, ng | PCR kit | Cycles post-PCR | output DNA lib |
| --- | --- | --- | --- | --- | --- | --- |
| 1A | RSMU_exome | Agilent v6 | 4800 | KAPA | 7 (½ volume) | 139 ng |
| 1B | RSMU_exome | Agilent v6 | 4800 | KAPA | 7 (½ volume) | 144 ng |
| 2A | RSMU_exome | MGI v4 | 4800 | KAPA | 10 (½ volume) | 520 ng |
| 2B | RSMU_exome | MGI v4 | 4800 | KAPA | 10 (½ volume) | 585 ng |
| 3A | MGI | MGI v4 | 2000 | MGI | 13 | 127 ng |
| 3B | MGI | MGI v4 | 2000 | MGI | 13 | 123 ng |

### Probe design comparison

Each manufacturer of the exome enrichment kit tries to select the most successful probe design and select the most relevant areas of the human exome for targeted capture. The design of MGIEasy v4 and Agilent v6 probes have many similarities (Table 8). Both manufacturers use biotinylated cRNA probes to hybridize target DNA, with MGIEasy v4 probes being 30 b shorter than Agilent v6 probes. We measured probe concentrations with the Qubit RNA HS Assay system (Life Technologies), and Agilent v6 probes were 2-fold more concentrated than MGIEasy v4 probes.

**Table 89:** Exome capture bait designs.

|  | MGI v4 | Agilent v6 |
| --- | --- | --- |
| Bait type | biotinylated cRNA baits | biotinylated cRNA baits |
| Bait length range, bases | 90 | 120 |
| Bait concentration, | 114 | 218 |
| ng/mkL |  |  |
| Target size, Mb | 59 | 60 |
| Number of target regions | 198 025 | 243 872 |
| Median Region Size, bp | 210 | 210 |
| .bed file (link) | MGIEasy Exome Capture V4 Probe Set<br>https://en.mgi-tech.com/products/reagents_info/id/9 | S07604514_Covered.bed<br>https://earray.chem.agilent.com/suredesign/index.htm, SureSelect Human All Exon V6 r2, design identification number S07604514 |

The MGIEasy v4 set targets 198,025 regions of the human genome of size 59 MB, the Agilent v6 set targets a slightly larger number, 243,872 regions of size 60 MB. Median Region Size (bp) is the same for both platforms = 210 bp. We compared bed files for MGIEasy v4, Agilent v6 to determine the degree of target region overlap between these two probe sets. We also looked at the coverage of target regions using the ENSEMBL protein coding exon database (Ensembl bed file for GRCh37/hg19 assembly, ensembl genes track for coding exons was obtained from https://genome.ucsc.edu/cgi-bin/hgTables). To illustrate the overlap of the target regions of the MGI v4 exome, Agilent v6 exome, and Ensembl coding exons, we draw Venn diagrams with their target size, using the matplotlib-venn library (https://github.com/konstantint/matplotlib-venn) (Figure 3).

**Figure 3.**
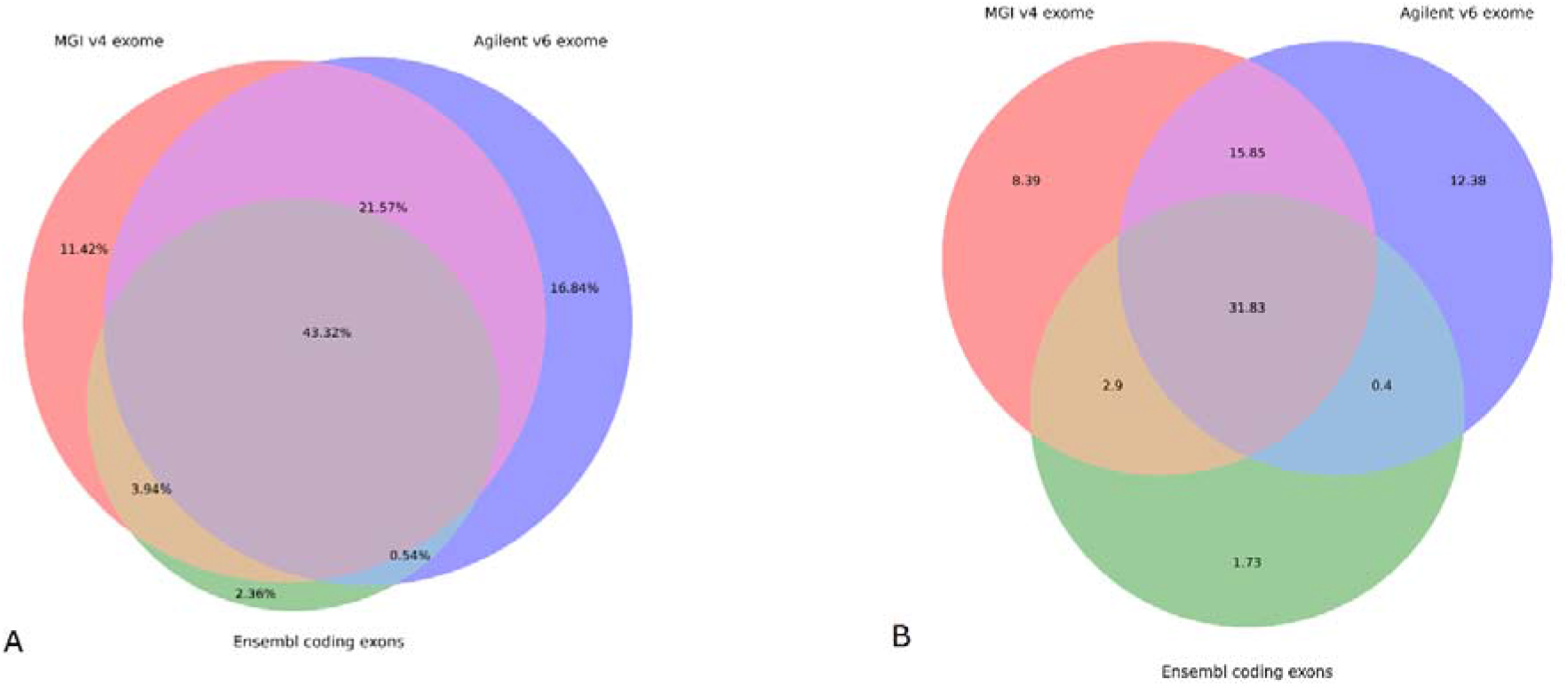
Weighted (in %) (A) and not-weighted (in Mb) (B) Venn diagram and for design intersection of MGI v4 exome, Agilent v6 exome, Ensembl coding exons. In Figure 3A depicts the intersection of all three bed files based on their target size with values of overlaps and unique regions as a percentage. Figure 3B shows a vein diagram for MGI v4 exome, Agilent v6 exome, and Ensembl coding exons without considering their real size, where values for different regions are in megabases (MB).

The percentage of unique target regions for the MGI v4 exome is 11.42% (8.39 MB), for the Agilent v6 exome is 16.84% (12.38 MB) and for Ensembl coding exons it is 2.36% (1.73 MB). The overlap area of all three bed files is 43.32% (31.83 MB). The overlap area between Agilent v6 and MGI v4 exome is 64.89% (47 MB). Meanwhile, the overlap between MGI v4 and Ensembl (47.26%) is greater than the overlap between Agilent v6 and Ensembl (43.86%). The percentage of uncovered Ensembl coding sites by either set was 2.36%.

### Raw data and Pooling balance

For each pool, about 324,5-434,5 millions (M) of aligned 150-bp paired-ends reads were generated in two MGISEQ-2000 runs.

The results of the fastQC quality check are presented as a single report collected by multiqQC [21], as well as separate reports for each sample (S2 File). Most data are Phred+35 (average quality per read) which indicates high sequence quality. Based on the results of FastQC reports, we often observe slight base imbalance in the Per base sequence content parameter at the beginning of reads and very rarely at the end, which are usually caused by non-random fragmentation and processing of the ends of DNA molecules during library preparation. We evaluated the level of imbalance and, if necessary, trimmed several bases, usually from 1 to 3 bases at the beginning of the read and 0-2 at the end.

It should be noted that the distribution of Per sequence GC content in the fastQC reports for samples enriched with Agilent v6 probes has a bimodal structure. Interestingly, the GC content distribution of exons in the human genome is also bimodal in shape [27, 28], so the Agilent solution for this parameter looks like it is closer to the regions of interest in the genome.

We obtained an average 65-75 M reads per sample, but within pools for different protocols the variance in the number of reads per sample was: pools of 12 samples - 1A - Δ53 M reads, 1B - Δ35 M reads, 2A - Δ22 M reads, 2B - Δ23 M reads; pools of 8 samples - 3A - Δ64 M reads, 3B - Δ101 M reads. Figure 4 shows a stacked barplot with the distribution of MB across samples in pools. Also, for clarity, we plotted the quantile function (Figure 5), where we can see the dynamics of data size distribution. It is seen that the MGI protocol and MGI v4 exome probes, represented by the green line (pools 3A + 3B), look unbalanced compared to the other pools (Figure 5A). The sharp jump at the beginning of the quantile function, indicates that the proportion of samples receiving less than 12,000 megabases is 0.4. In Figure 5B, we see a more detailed picture of the imbalance between pools 3A and 3B. Such specificity of data acquisition does not allow achieving an even distribution of MB over the samples, which may entail under-coverage of some regions in the samples in the pool. Although in pools 1A+1B we see a slight imbalance at the very beginning, the balance of the obtained sample data is higher for the RSMU_exome protocol than for MGIEasy protocol.

**Figure 4.**
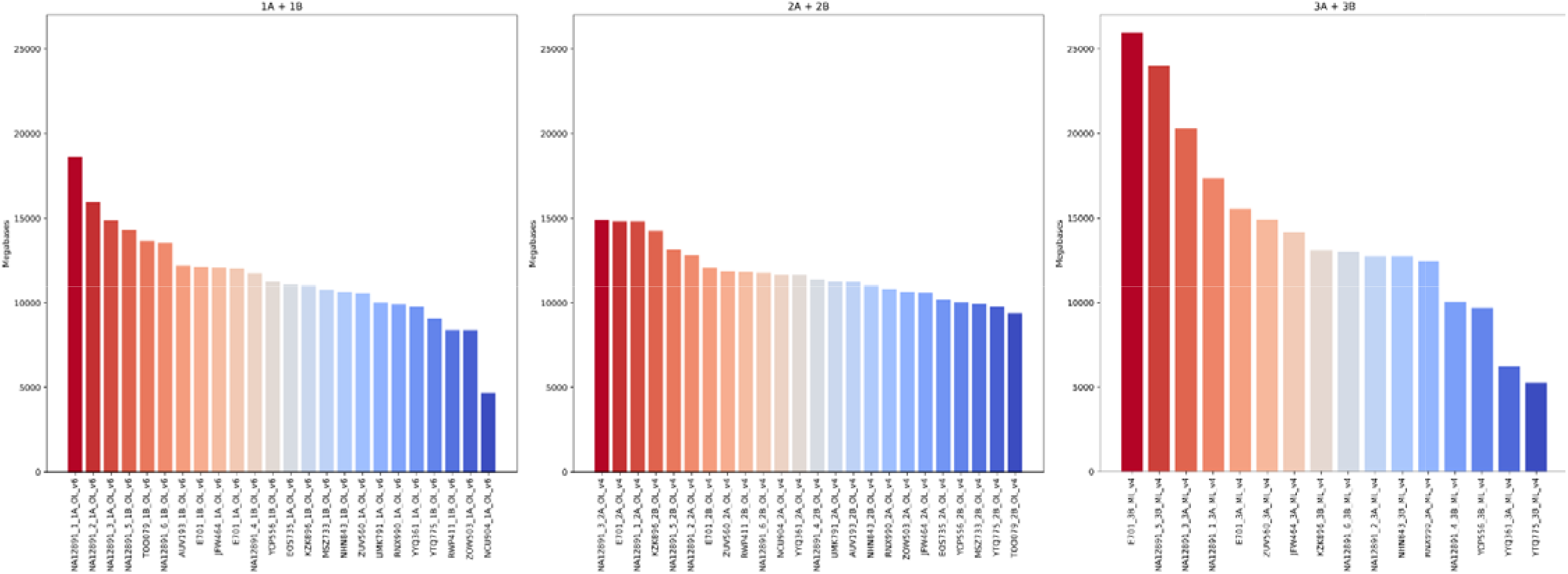
Stacked barplot for sample sizes by pool in megabases.

**Figure 5.**
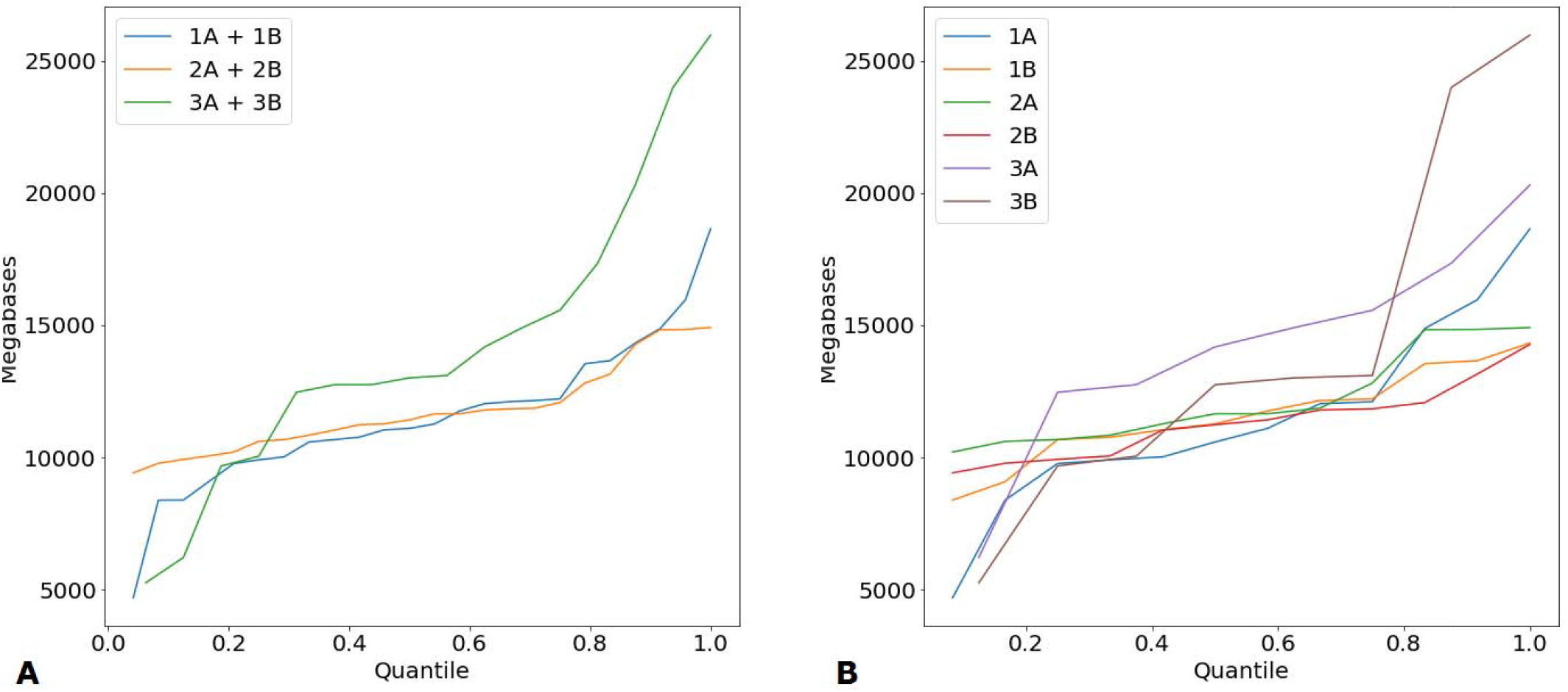
(A) Quantile function for averaged pools 1A+2A, 1B+2B, and 3A+3B. (B) Quantile function for each of the pools.

### Quality of enrichment

At least 99% of the reads for each sample were mapped to human DNA using BWA. For all samples in the pools, we calculated standard metrics for on-target %, off-target %, mean coverage, % of regions covered at x5, x10, x20, and others (S3 Table) using Picard. For pools 1A and 1B we got 6.3-15.4% (mean 7.3%) of duplicates, 2A and 2B 15.2-22.5% (mean 12.4%), 3A and 3B 10-26.5% (mean 11.4%), the more reads, the more duplicates. The minimum value of the “Covered x10” parameter for the samples with the lowest number of reads for approach 1 was 96,87 %, for approach 2 - 96,12 %, for approach 3 - 86,56 %; The lowest mean coverage of samples in pools 1A and 1B = 69.1 (52.39 M reeds) and 67.73 (52.37 M reeds), in pools 2A and 2B 79.82 (58.03 M reeds) and 74.26 (54.48 M reeds), in pools 3A and 3B 43.81 (35.92 M reeds) and 30.9 (31, 57 M reeds) respectively.

### Comparison of methods with downsampled Platinum Genomes

#### Mapping: on target percentage and duplication rate

To correctly compare the 3 different approaches, we downsampled the raw melon raw data to 50 million reads for Platinum Genomes samples using Picard v2.22.4 (S4 Table). We then estimated the proportion of on-target, off-target, un-aligned, and percentage of duplicates for each of the pools (Figure 6). For approach 1 - RSMU_exome protocol with Agilent v6 probes, the amount of duplicates is the lowest - 5.8%, while approaches 2 and 3 made with RSMU_exome and MGIeasy protocols with MGI v4 probes have the duplicate amount of about 10%. For all approaches, the number of un-aligned reads does not exceed 1%. The percentage of masked off-target reads was 7.6% for approach 1 (RSMU_exome protocol + Agilent v6), 11% for approach 2 (RSMU_exome protocol + MGI v4), and 19% for approach 3 (MGIeasy protocol + MGI v4).

**Figure 6.**
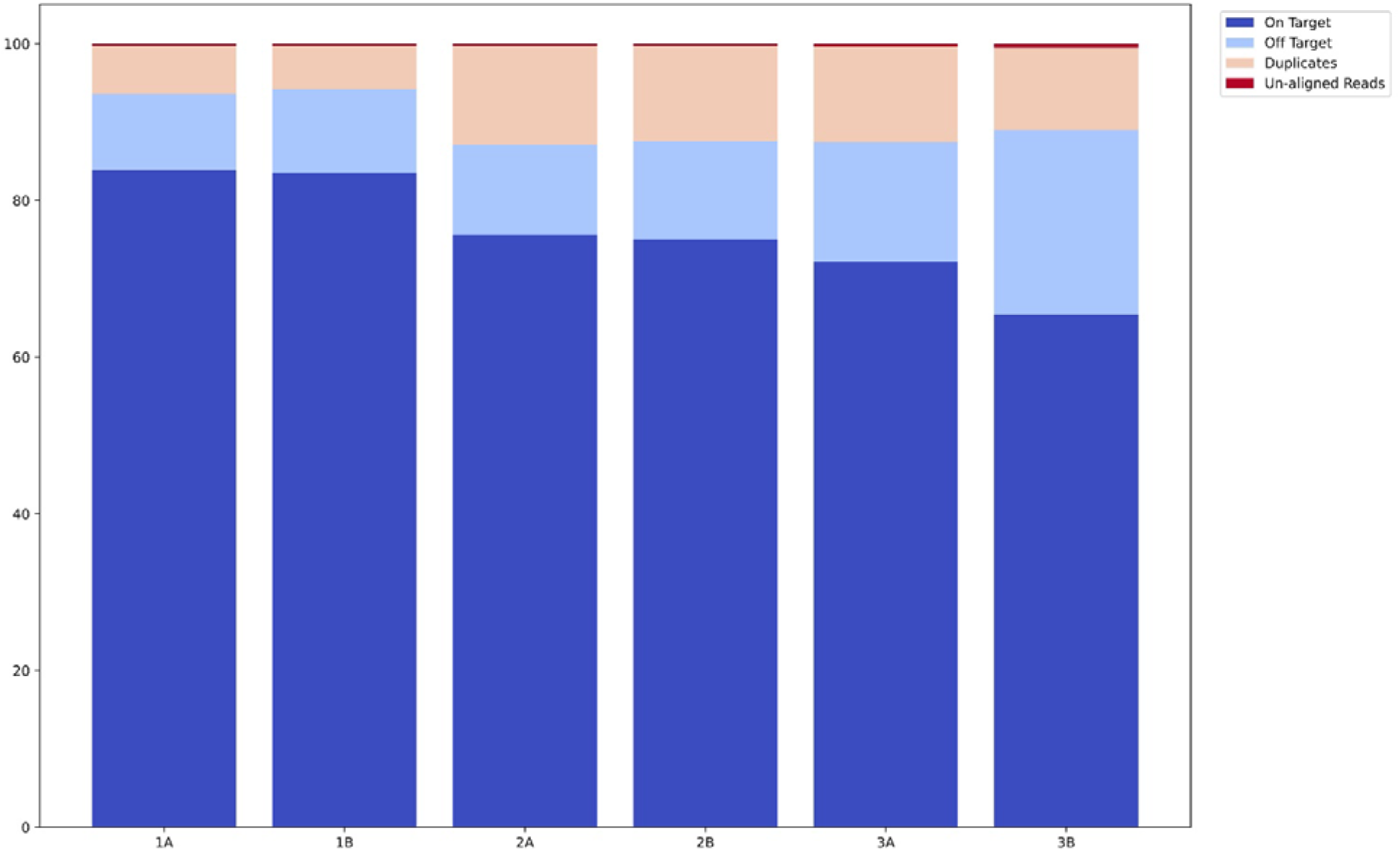
Stacked barplots for on-target, off-target, duplicates, and un-aligned reads averaged within pools.

### Coverage analysis

Enrichment efficiency was assessed by comparing the depth of coverage of target regions for the 3 approaches. Only NA12891 samples downsampled to 50 M and, if necessary, to 40 M, 30 M, and 20 M reads participated in the pool coverage comparison. Coverage values for samples within pools A and B were averaged because we see no significant differences between the results for pools A and B within each approach, indicating a good level of technical reproducibility. Coverage was evaluated on 3 bed files: bed file corresponding to the probes on which the samples were made, bed file for ensembl coding exons, bed file of shared regions for MGIEasy v4 and Agilent v6 (cross bed). See S4 Table for Picard data for all samples downsampled to 20 M, 30 M, 40 M, and 50 M reads.

The fraction of target regions covered at least 1 time for the compared approaches differed slightly on downsampled to 50 M samples (Figure 7A), with the MGI probes averaging almost the same for both our protocol (98.60%) and the original MGIeasy protocol (98, 59%); for Agilent v6 probes the value was 97.93%. At the same time, the trend changes markedly for “on-target covered x10”, where our approach RSMU_exome with Agilent v6 probes maintains coverage at a high level, while the MGIeasy protocol loses dramatically in values. Thus, the average “on-target covered x10” were: 95,81% (min = 94.25%) for RSMU_exome protocol + Agilent v6; 94.96% (min = 94.4%) for RSMU_exome protocol + MGI v4; 91,13% (min = 88.15%) for MGIeasy protocol + MGI v4. The parameter “on-target covered x30” is on average > 75% for the RSMU_exome, and lower by 12% for the MGIeasy (on average > 63%).

For bed ensemble coding exons (target size= 35.4 Mb) and for overlaped target regions MGI v4 and Agilent v6 (cross bed, target size = 47.9 Mb) the trend remained, the RSMU_exome approach on both bed files was better.

**Figure 7.**
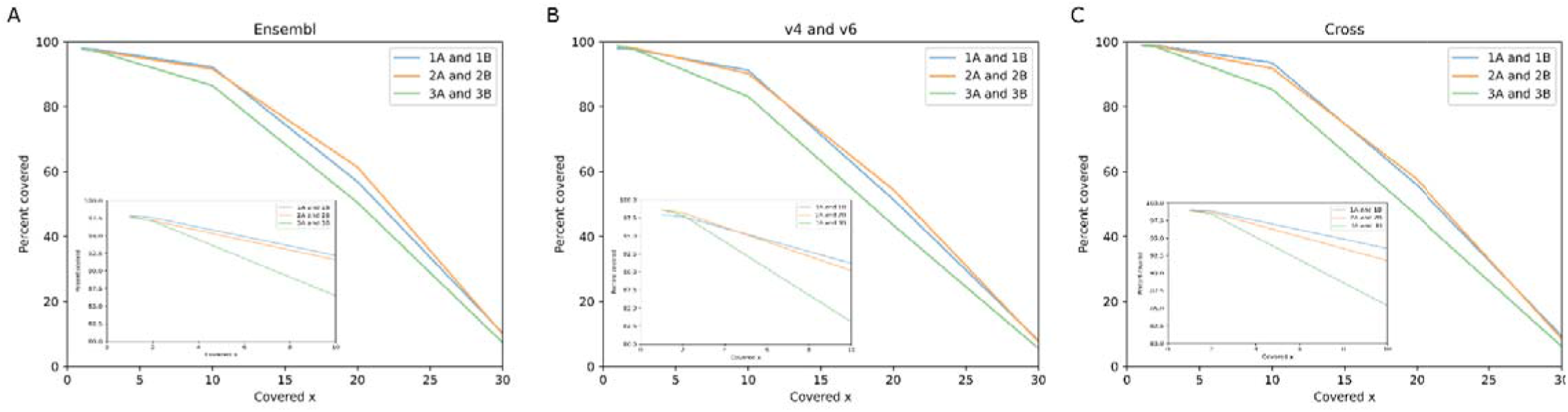
Dependence of region coverage quality for different depths on 50 M read samples for regions corresponding to bed files: (A) Ensembl, (B) - sample probes (MGI v4 or Agilent v6), (C) - intersection regions of bed files MGI v4 and Agilent v6.

We also examined the % on-target covered x10 for downsampled samples up to 10M, 20M, 30M, 40M, and 50M reeds (Figure 8). The RSMU_exome protocol shows the best results for all beds - already 30M reads on average are sufficient to cover x10 over 90% of the on-target regions on Agilent v6, MGI v4, and cross bed, and over 91% for ENSEMBL coding regions. For the MGIEasy protocol, such results are achieved from 50 M reads/sample. The curves plateaued faster with less than ~2% on- target coverage at ≥10× for the RSMU_exome protocol for both Agilent v6 and MGI v4 probes at 40 M reads, in contrast to the MGIeasy protocol for MGI v4 probes, indicating higher hybridization and capture efficiency with our protocol. Thus, fewer reads are required to obtain the same completeness of exome coverage when RSMU_exome protocol is used.

**Figure 8.**
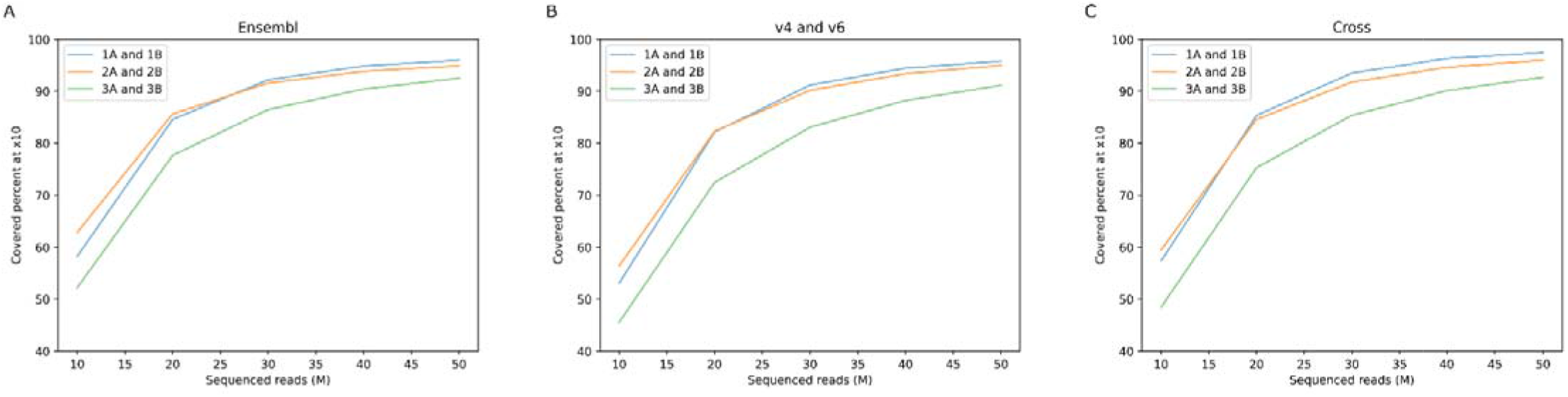
Percentage of regions covered x10 for downsampled samples for corresponding bed files: (A) ensembl, (B) sample probes (MGI v4 or Agilent v6), (C) overlapping regions of bed files MGI v4 and Agilent v6.

### GC content

Most enrichment techniques show fall read depth in GC-rich and GC-poor DNA regions, and we compared pools on this trait using the density plots (Figure 9). Expectedly, all 3 approaches showed a drop in depth coverage in extremely low (<20%) and high (>80%) GC content regions. Interestingly, pools 2A+2B and 3A+3B enriched in different protocols but with MGI v4 probes have more even coverage between 40% and 60% GC, in contrast to pools 1A+1B enriched with Agilent v6 probes, for which the overcoverage peak (>x400) of target regions shifted towards 50% and 70% GC. For pools 1A+1B the highest density of coverage is in the region of 40%-50% GC. There is little difference between pools 2A+2B and 3A+3B (MGI v4 probes), although the coverage density between 40% and 60% GC is more uniform for pools 2A+2B made using the RSMU_exome protocol.

**Figure 9.**
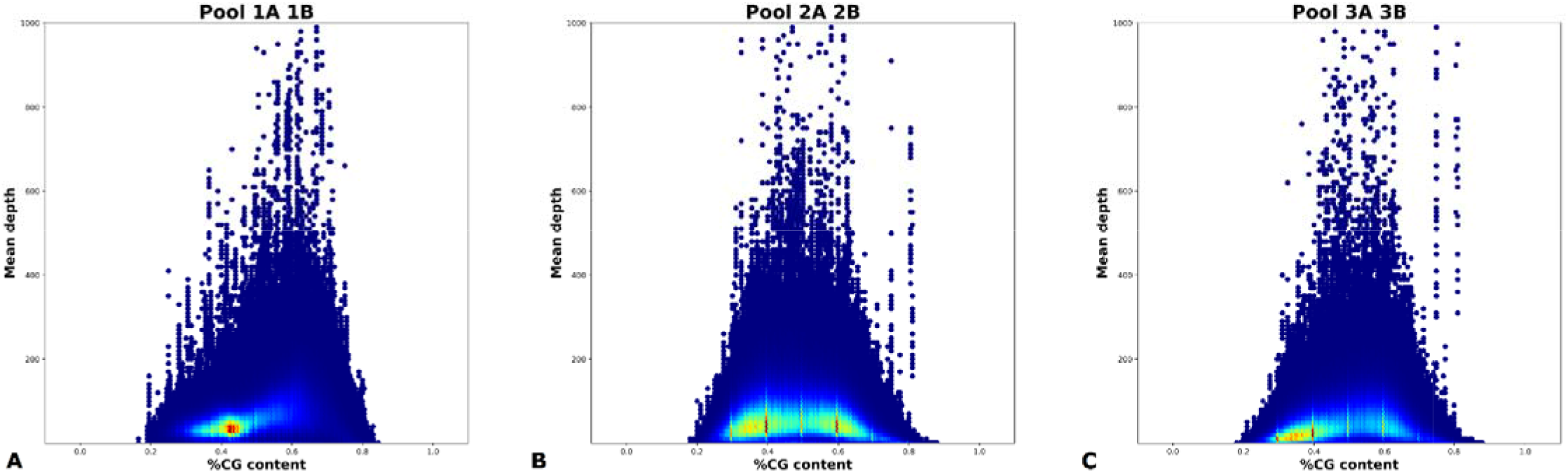
Density plot of %GC Content vs Mean Depth for A) RSMU_exome + Agilent v6; B) RSMU_exome + MGI v4; C) MGIeasy capture + MGI v4. Here we show 2D density plot of %GC Content vs Mean Depth parameters which were collected by Picard HsMetrics v2.22.4. We collected data for this plot by merging all samples from corresponding pools (1A + 1B, 2A + 2B, 3A + 3B) by Picard 2.22.4. Density estimation was performed using 2D histograms. More specifically we chose data points in fixed rectangle (%GC Content □[0;1] and Mean Depth □[0;1000]), then we split this rectangle into evenly spaced grid of size 200×100 and counted how many data points lie in each cell in the grid. Finally we normalized grid to [0,1] range and plotted it using “jet” colormap from matplotlib library.

### SNV and CNV analysis

For SNV analysis, we used data filtered using the algorithm described above, including deduplication and downsampling to 50 M reads per sample. Next, using bcftools mpileup v1.9, we obtained information about gene sequence variations. We filtered them by coverage, leaving only the variants covered by at least 13x and those localized inside the target.

To assess the similarity of the data obtained, the IoU values between all samples from the 6 pools were calculated. In Figure 10 we can clearly see two clusters that form the samples NA12891 and E701. These clusters provide clear evidence that the data for NA12891 are replicates libraries, despite the fact that they were prepared using different protocols. Meanwhile, for replicates of NA12891 samples from pools A and B within each approach, the IoU values show maximum results, which shows excellent reproducibility of results for different pools for the same approach.

**Figure 10.**
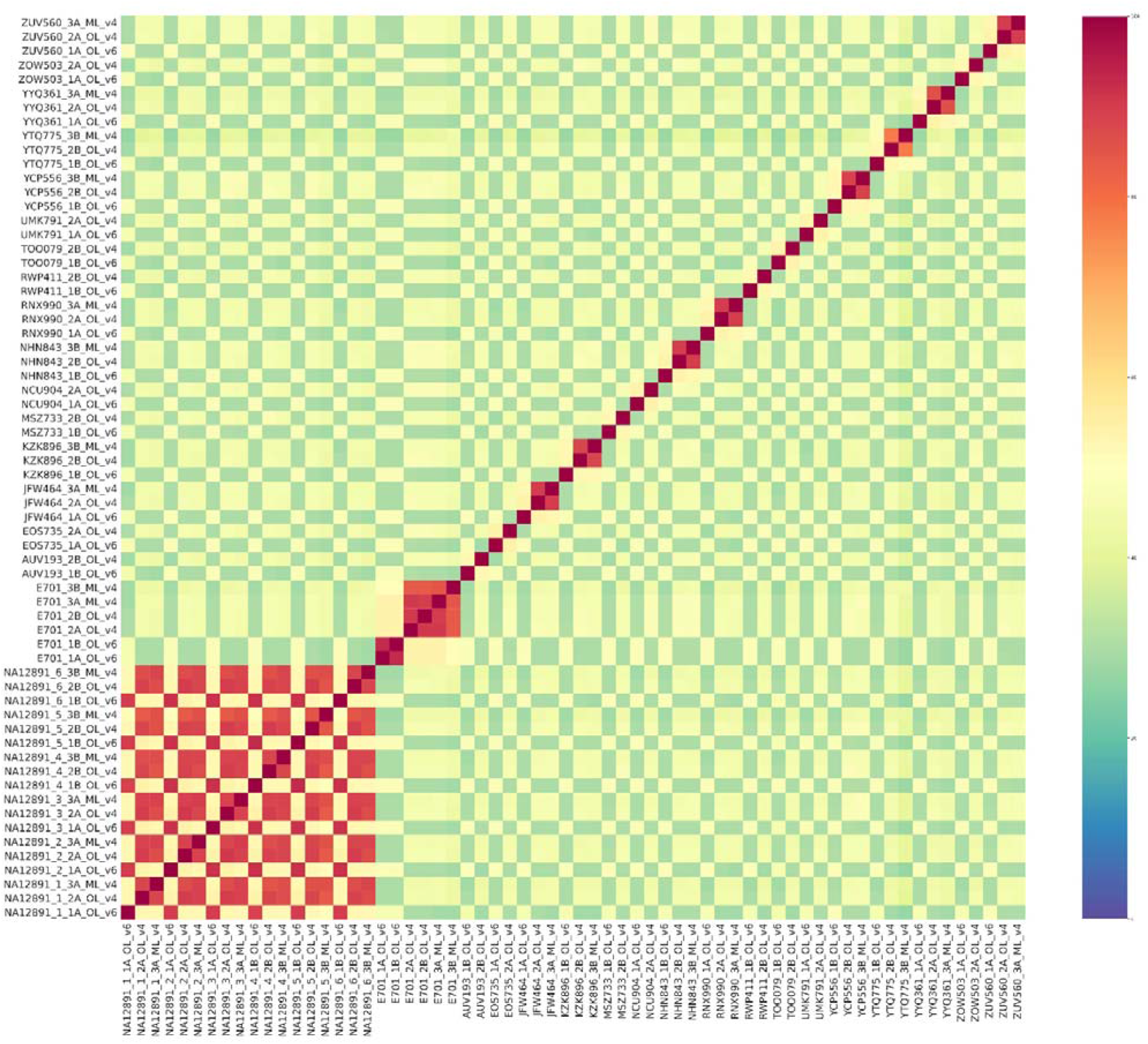
SNV Heatmap for all samples participating in the experiment. Visualization of IoU SNV results for all samples filtered by target regions.

To understand whether our results for the platinum genome are valid, we used NA12891 WGS data from the open source (ftp://ftp.sra.ebi.ac.uk/vol1/fastq/SRR622/SRR622458) and using IoU, considering position, type of substitution (SNV or CNV) and genotype, we evaluated the overlap of one of our NA12891 samples into the downloaded reference genome, which was also counted according to our bioinformatic pipeline. We used a Venn diagram to visualize this - Figure 11 shows that for samples filtered by Agilent v6 bed file, our NA12891 sample from pool 1A almost completely fits into the reference genome NA12891_ref37 with a depth-of-read cutoff over 13 reads. From the resulting Venn diagram, we can conclude that the results for our samples are correct, and bcftools mpileup v1.9 allows correct variant calling with fairly good accuracy.

**Figure 11.**
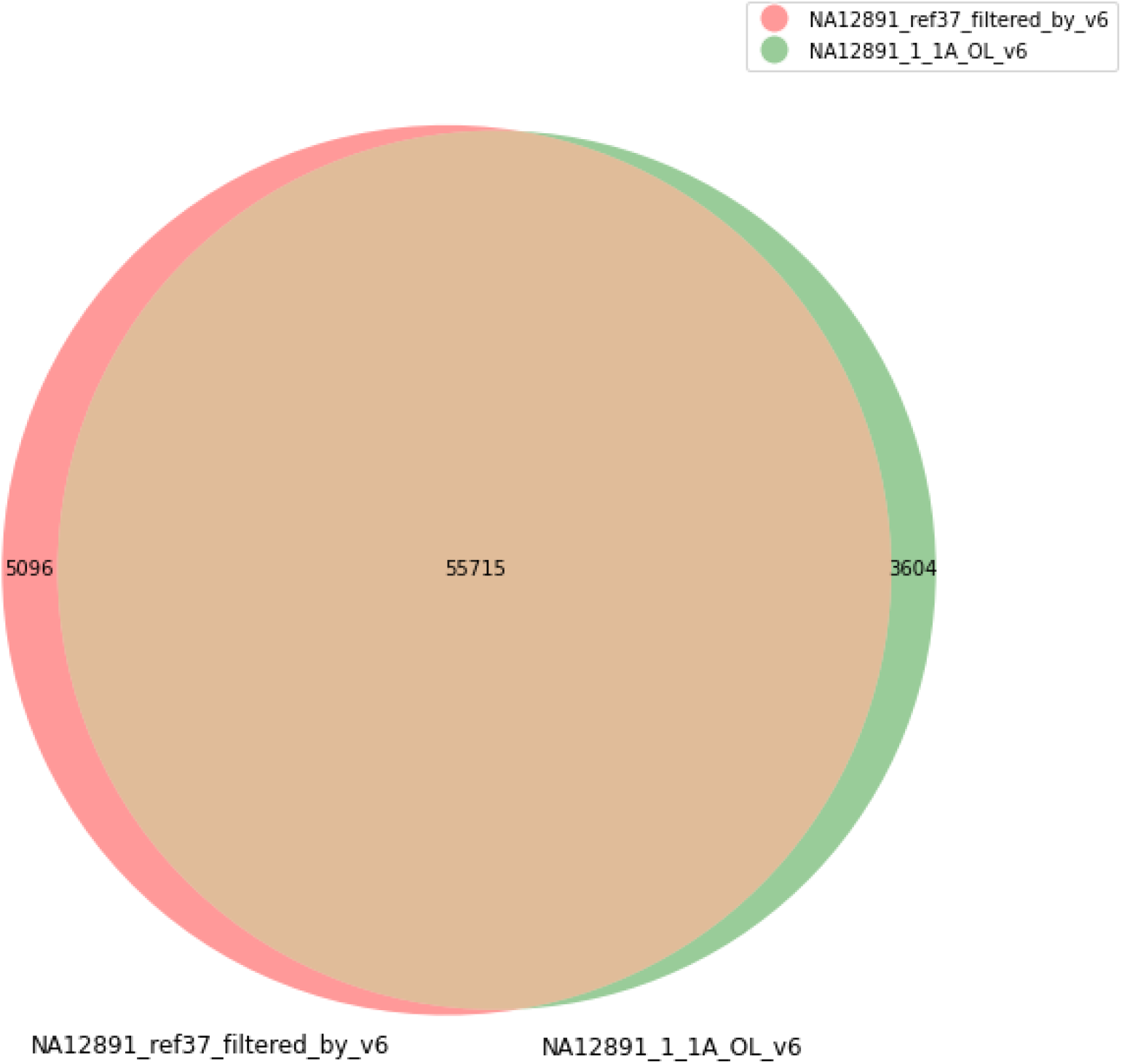
Venn diagram for the intersection of filtered SNV and CNV for randomly chosen sample NA12891 from pool 1A and the reference genome of sample NA12891. The data were filtered for depth of coverage over 13 reads. The 55,715 SVNs and CNVs found were a complete match to the reference genome by genotype.

To estimate the quality of CNV calling with bcftools mpileup v1.9 on our data pre-filtered on target regions, we used the IoU metric. We calculated Intersection over Union values only for lines from vcf that contained insertions and deletions. Based on the results, we constructed two heatmaps Figure 12 for indels and deletion separately. A random sample E701 from pool 1A was added for clarity.

**Figure 12.**
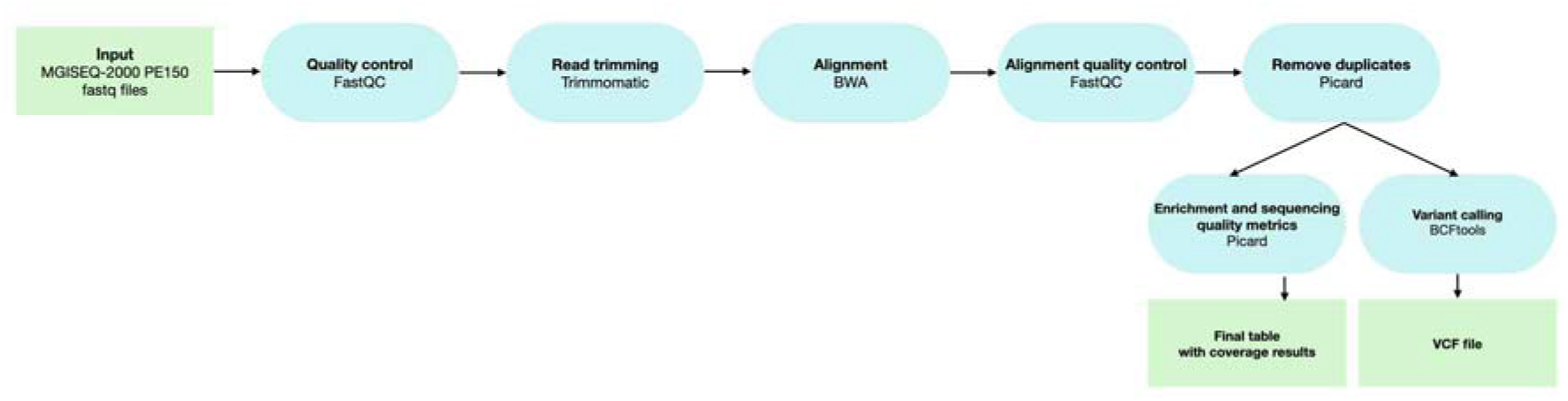
Schematic of the bioinformatic pjpeline for exome data processing. The figure shows the programs that were applied to analyze the data obtained both during the experiment and downloaded from free sources.

As we can see on the heatmap, calling copy number variation with bcftools mpileup v1.9 allows better calling deletion than insertions. Samples from different pools have less similarity than samples within pools, but despite this, we can see a significant difference between the CNV for sample NA12891 and sample E701.

**Figure 12.**
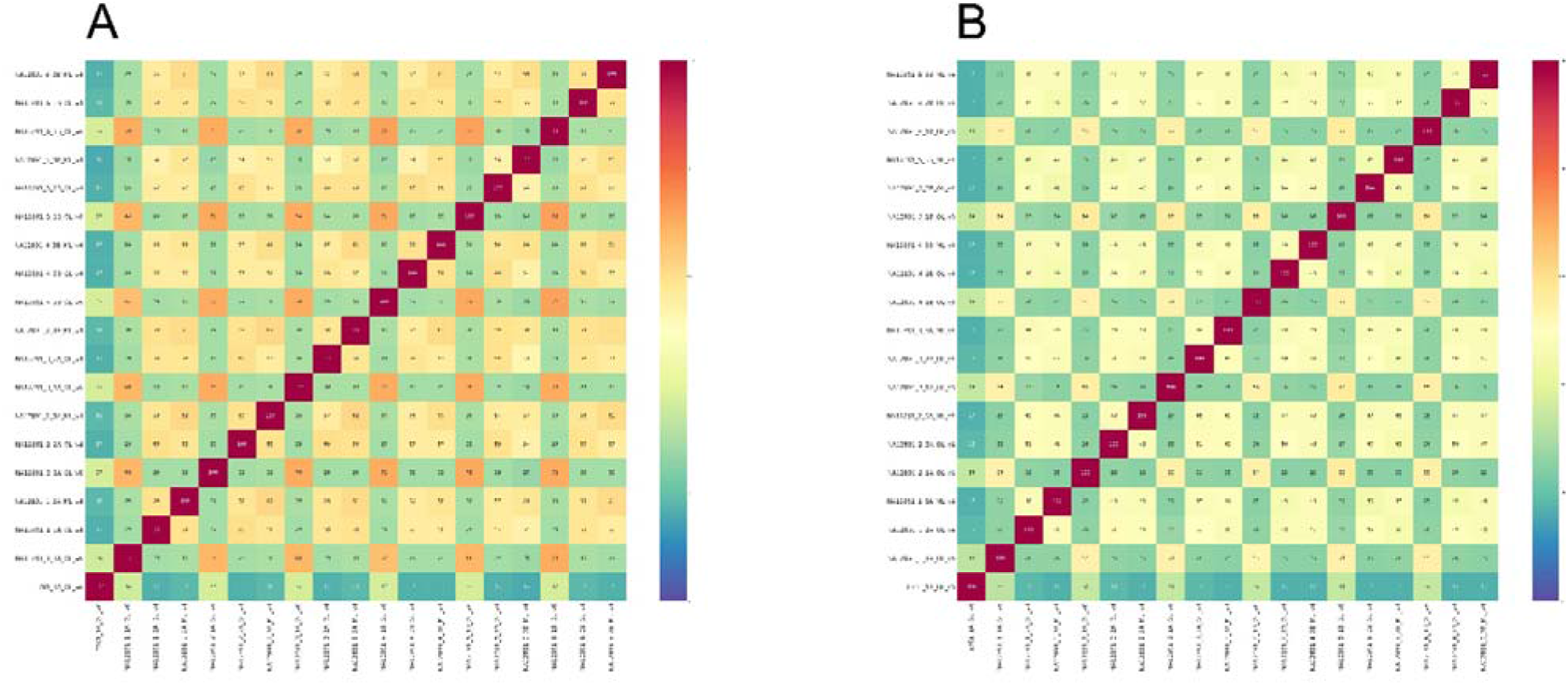
IoU hitmaps for CNVs from NA12891 samples and one random sample E701 from pool 1A. Visualization of IoU results for insertions (A) and deletions (B), which can be called using bcftools mpileup v1.9.

Also, we collected pool statistics for variant calling data. All data from the pools were sorted by their respective bed file and by target regions for ensembl coding exons. In this estimation method, we looked at the parameters of the number of total SNVs and CNVs and those that hit only the target regions.

The mean total number of SNVs in on-target regions was 57 189, (RSMU_exome + Agilent v6), 54 094 (RSMU_exome + MGI v4), 54 044 (MGIeasy protocol + MGI v4) when using the 50M read sets in the NA12891 sample.

We varied the filtering parameters of the variants, the results are presented in S5 Table. We noticed that the QUAL>20 and QUAL>30 parameters give a large cutoff on the entire number of base substitutions, but with almost no change in the values for the results within the target regions. The strictest cutoff for variants with depth of coverage greater than 13 reads (DP > 13) and with the QUAL>30 parameter had little effect on the number of variants. For example, the values for SNV for pool 1A changed by 3.1% (from 57,189 to 55,407), which is not a critical change. Also, we noticed that pools 1A and 1B had higher values for the absolute number of all SNVs and CNVs from full vcf file and those in target regions. With the strictest SNV cutoff (DP > 13, QUAL>30), the number of variants in the target for pools 1A and 1B averages 55 thousand variants, for pools 2A and 2B 50 thousand, and for pools 3A and 3B about 49 thousand. Approach 1 shows the best results. At the same time, in the case of a fixed probe set (MGIEasy v4) and differences only in the enrichment process, with the same sequencing data volume our RSMU_exome protocol allows to obtain 2% more qualitative data for variant calling than the MGIEasy protocol.

In general, we can note that for all cutoffs, the average SNV and CNV for samples made on Agilent v6 within their target regions is higher than for MGI v4. But, in the case of ensembl genes track for coding exons, we do not see strong differences between the pools.

### Statistical analysis

To assess the quality of SNV and CNV detection, we took the variant calling results of the platinum genome libraries in all 6 pools. Samples were pre-filtered by target regions for the corresponding set.

When evaluating the quality of SNV and CNV detection, we used platinum genome data as a reference. The reference data were pre-filtered by target regions (MGI v4, Agilent v6), according to the type of probes used on which our sample was made.

**Table 6.** Variation accuracy estimation by comparison with reference Platinum Genome for SNV (PG).

| Filename | Total | In PG | Not in PG | PG-specific variations | SNV Sensitivity | SNV Precision | SNV F-measure |
| --- | --- | --- | --- | --- | --- | --- | --- |
| NA12891_2_1A_OL_v6.vcf | 58951 | 56252 | 2699 | 3267 | 99.652% | 99.905% | 99.778% |
| NA12891_5_3B_ML_v4.vcf | 51568 | 49202 | 2366 | 7976 | 99.667% | 99.926% | 99.796% |
| NA12891_4_3B_ML_v4.vcf | 53305 | 50801 | 2504 | 6377 | 99.518% | 99.931% | 99.724% |
| NA12891_3_1A_OL_v6.vcf | 58984 | 56290 | 2694 | 3229 | 99.561% | 99.906% | 99.733% |
| NA12891_1_2A_OL_v4.vcf | 55359 | 52431 | 2928 | 4747 | 99.549% | 99.935% | 99.742% |
| NA12891_1_1A_OL_v6.vcf | 57867 | 55124 | 2743 | 4395 | 99.609% | 100.000% | 99.804% |
| NA12891_6_3B_ML_v4.vcf | 52974 | 50381 | 2593 | 6797 | 99.656% | 100.000% | 99.828% |
| NA12891_3_2A_OL_v4.vcf | 55510 | 52764 | 2746 | 4414 | 99.558% | 99.968% | 99.762% |
| NA12891_2_2A_OL_v4.vcf | 55589 | 52698 | 2891 | 4480 | 99.553% | 99.968% | 99.760% |
| NA12891_5_2B_OL_v4.vcf | 55010 | 52305 | 2705 | 4873 | 99.680% | 99.904% | 99.791% |
| NA12891_1_3A_ML_v4.vcf | 53619 | 50883 | 2736 | 6295 | 99.658% | 99.863% | 99.761% |
| NA12891_6_1B_OL_v6.vcf | 58371 | 55746 | 2625 | 3773 | 99.648% | 99.936% | 99.792% |
| NA12891_4_2B_OL_v4.vcf | 54597 | 51903 | 2694 | 5275 | 99.704% | 100.000% | 99.852% |
| NA12891_4_1B_OL_v6.vcf | 59013 | 56378 | 2635 | 3141 | 99.533% | 99.938% | 99.735% |
| NA12891_3_3A_ML_v4.vcf | 54230 | 51625 | 2605 | 5553 | 99.633% | 99.933% | 99.783% |
| NA12891_6_2B_OL_v4.vcf | 54807 | 52063 | 2744 | 5115 | 99.676% | 100.000% | 99.838% |
| NA12891_2_3A_ML_v4.vcf | 54143 | 51421 | 2722 | 5757 | 99.627% | 99.966% | 99.796% |
| NA12891_5_1B_OL_v6.vcf | 59109 | 56439 | 2670 | 3080 | 99.626% | 99.875% | 99.750% |
| **Total** | 1003006 | 954706 | 48300 | 88544 | 99.552% | 99.765% | 99.658% |

**Table 7.** Variation accuracy estimation by comparison with reference Platinum Genome for CNV (PG).

| Filename | Total | In PG | Not in PG | PG-specific variations | CNV Sensitivity | CNV Precision | CNV F-measure |
| --- | --- | --- | --- | --- | --- | --- | --- |
| NA12891_2_1A_OL_v6.vcf | 58951 | 56252 | 2699 | 3267 | 99.994% | 99.979% | 99.987% |
| NA12891_5_3B_ML_v4.vcf | 51568 | 49202 | 2366 | 7976 | 99.996% | 99.981% | 99.988% |
| NA12891_4_3B_ML_v4.vcf | 53305 | 50801 | 2504 | 6377 | 99.996% | 99.971% | 99.983% |
| NA12891_3_1A_OL_v6.vcf | 58984 | 56290 | 2694 | 3229 | 99.994% | 99.974% | 99.984% |
| NA12891_1_2A_OL_v4.vcf | 55359 | 52431 | 2928 | 4747 | 99.996% | 99.972% | 99.984% |
| NA12891_1_1A_OL_v6.vcf | 57867 | 55124 | 2743 | 4395 | 100.000% | 99.977% | 99.988% |
| NA12891_6_3B_ML_v4.vcf | 52974 | 50381 | 2593 | 6797 | 100.000% | 99.979% | 99.989% |
| NA12891_3_2A_OL_v4.vcf | 55510 | 52764 | 2746 | 4414 | 99.998% | 99.972% | 99.985% |
| NA12891_2_2A_OL_v4.vcf | 55589 | 52698 | 2891 | 4480 | 99.998% | 99.972% | 99.985% |
| NA12891_5_2B_OL_v4.vcf | 55010 | 52305 | 2705 | 4873 | 99.994% | 99.980% | 99.987% |
| NA12891_1_3A_ML_v4.vcf | 53619 | 50883 | 2736 | 6295 | 99.992% | 99.979% | 99.985% |
| NA12891_6_1B_OL_v6.vcf | 58371 | 55746 | 2625 | 3773 | 99.996% | 99.979% | 99.988% |
| NA12891_4_2B_OL_v4.vcf | 54597 | 51903 | 2694 | 5275 | 100.000% | 99.982% | 99.991% |
| NA12891_4_1B_OL_v6.vcf | 59013 | 56378 | 2635 | 3141 | 99.996% | 99.972% | 99.984% |
| NA12891_3_3A_ML_v4.vcf | 54230 | 51625 | 2605 | 5553 | 99.996% | 99.977% | 99.987% |
| NA12891_6_2B_OL_v4.vcf | 54807 | 52063 | 2744 | 5115 | 100.000% | 99.980% | 99.990% |
| NA12891_2_3A_ML_v4.vcf | 54143 | 51421 | 2722 | 5757 | 99.998% | 99.977% | 99.988% |
| NA12891_5_1B_OL_v6.vcf | 59109 | 56439 | 2670 | 3080 | 99.992% | 99.977% | 99.985% |
| Total | 1003006 | 954706 | 48300 | 88544 | 99.985% | 99.972% | 99.979% |

Sensitivity, precision, and F-measure metrics were used to assess the quality of SNV and CNV detection. Sensitivity estimates the probability that if our variant calling method detects a position in the genome as SNV/CNV, then it is in fact SNV/CNV. Precision estimates the likelihood that if a position in the genome is actually SNV/CNV, then our method will detect it as SNV/CNV. These two metrics are indicative of the quality of the method. We also calculated the aggregation metric F-measure, which is the average of harmonic sensitivity and precision. It is worth noting that the detection of SNV and CNV is a task of binary classification into two categories (respectively SNV and CNV). In this regard, the metrics for SNV and CNV cannot be considered independent, because they are derived from the same metric matrix.

As a result, we see that SNV and CNV detection quality metrics show excellent results (Table 6, Table 7). It is also worth noting that, in general, the number of SNV and CNV in the nutrient target of samples from pools 1A and 1B prepared according to our protocol shows slightly higher results. Based on these results, we can conclude that the data are of high quality, which allows us to detect the maximum number of SNV and CNV.

## Discussion (Belova, Pavlova, Afasizhev)

To perform high-quality WES analysis for bulk libraries in the laboratory on the MGISEQ-2000 sequencer and after testing the MGIEasy v4 enrichment protocol and being dissatisfied with the results, we decided to develop our enrichment protocol. Having studied the protocols of commonly known exome enrichment kits and compared them [29,30,31,32], we focused on obtaining high quality enrichment of each multiplexed sample with minimal sequencing.

To validate the protocol, we used the platinum genome NA12891. To minimize PCR errors and the number of duplicates, we reduced the number of post-capture PCR cycles to 7 cycles by introducing the hybridization and capture protocol modifications described above. One of the peculiarities of sample preparation for sequencing on the MGISEQ-2000 is the stage of circularization of prepared libraries, which, in our experience, requires at least 80-100 ng of libraries. Accordingly, the yield after enrichment should be sufficient for further processing of the pool with a minimum number of PCR cycles, which corresponded to our protocol.

The maximum number of libraries per pool was increased to 12, which is more than in the MGIEasy Exome and Agilent SureSelect protocols (8 samples per pool), with increasing the amount of each DNA library to 400 ng per pool. It was shown that keeping 500 ng per sample per pool, regardless of the total amount of DNA entering the enrichment, the number of duplicates did not increase and sample coverage remained uniform [33]. We did not add Salmon Sperm DNA to the enrichment reaction because we assumed that it could hybridize with probes targeting conserved regions of the human genome, reducing the efficiency of the reaction.

Obviously, pre-capture multiplexing makes the enrichment procedure cheaper (~2.5-5 times) and also reduces the amount of time spent on the procedure, especially when the laboratory has a low level of process robotization [33-36]. However, obtaining an equal amount of data per sample after multiplex enrichment is not an easy task. In many studies, pooling eveness is described rather streamlined or not discussed at all, and after sequencing the difference in sample coverage in the pool can be as much as 10 times [9,34,35,37]. In [35], the authors pooled up to 16 samples before enrichment using the Agilent SureSelect XT protocol without loss of enrichment quality, but the difference in the number of M reads per sample can be as much as ~16 times. Other authors obtained a 10-fold difference in sample pool coverage for the Nextera kit [9]. The authors from the Center for Inherited Disease, show that in case of pre-capture pooling, the difference in the number of data/sample pooling can reach ~5 times for IDT protocol, ~2-3 times for Roche, Twist protocols [34]. Thus, the main disadvantage of pre-capture multiplexing is that samples “dropping out” of the pool require not just resequencing, but re-enrichment, which increases the final cost of analysis. According to the MGI enrichment protocol, the difference in the amount of data per sample reached x4 and 3 samples out of 16 failed to pass the threshold of on-target regions covered by 10x greater than 95% (samples NA12891_4_3B=94.73%, YYQ361_3A = 91, 46%, YTQ775_3B = 86, 57%), meaning they would have to be re-enriched. By the RSMU_exome protocol, there was less difference in the amount of data per sample, and no sample “dropped out.” The minimum value for 48 samples in the pools enriched with both Agilent and MGI on-target probes covered x10 = 96, 09%, which is an advantage of our technique.

After a correct comparison of samples with normalized coverage for each method, we were satisfied with the results obtained using the own RSMU_exome protocol for Agilent v6 and MGI v4 probes, both technologies yielded over 80-90% of bases on-target, although for the latter we obtained on average 2 times more duplicates. Meanwhile, Agilent probes, popular among researchers worldwide, showed slightly better on-target % and coverage of coding exons from the ENSEMBL database. However, per the results of other researchers [6-10], none of the probe designs was able to fully cover the sequences of all coding exons; regions that can escape not only WES, but also WGS [3,27] using short reeds are known, in particular, an increased number of genomic repeats or the presence of pseudogenes.

The specificities of sample preparation for WES can affect the quality of the variant calling even with the same target size [7]. Thus, when using the RSMU_exome protocol and the original MGIEasy both with MGI v4 probes, the calling results were better for the RSMU_exome protocol. Working with Agilent v6 probes using the RSMU_exome protocol the results of variant calling were comparable with the results of other researchers [3,6,7].

GC-content can affect the uniformity of coverage depth of target regions, i.e. low sequencing depth can be caused by high (>60%) or low (<40%) GC-content in the target. Such regions contribute to coverage due to reduced hybridization efficiency (which is affected by probe design) and postcapture PCR [38,39]. As expected, all 3 approaches showed bias for extreme GC regions, with the RSMU_exome protocol with MGI v4 probes being slightly more uniform between 40% and 60% GC-content. Our results for the Agilent SureSelect v6 probes are also similar to the results for the Agilent SureSelect QXT kit of the previous study by García-García et al [8] in their ability to cover GC-content regions.

Today for sequencing machines from MGI Tech there are few reagent kits for specific tasks, which allows manufacturers to set high prices for them. The presented modifications of the protocol of library preparation, pooling, and the protocol of enrichment and washing of libraries together turn out to be cheaper than the solution from MGI Tech by ~5 times per sample (sequencing is not included in the calculation).

We are the first to offer an alternative commercial protocol that is superior to the manufacturers’ off-the-shelf solutions in quality (high performance in terms of capture uniformity and on-target coverage with Agilent v6 probes, and with MGI v4 probes) and significantly cheaper than them, which we have demonstrated in this publication. We have posted the RSMU_exome protocol in its entirety in the S1 File so that other researchers can take advantage of it.

## Supplementary Materials

S1 File. «RSMU_exome» Protocol of the exome enrichment of library pools for WES sequencing on MGISEQ-2000

S2 File. MultiQC reports for sequenced exomes

S1 Figure. Bioanalyzer 2100 results for NA12891 libraries

S1 Table. Libraries summary information

S2 Table. Efficiency of PCR of post-capture pool of DNA libraries with or without Dynabeads C1

S3 Table. Picard metrics for all 64 samples

S4 Table. Picard metrics for downsampled NA12891 files

S5 Table. Variant calling results

## Author Contributions

**VB** – Conceptualization, Methodology, Investigation, Validation, Writing – Original Draft Preparation; **AP** and **RA** – Formal Analysis, Methodology, Software, Visualization, Writing – Original Draft; **VM, MK, VCh** – Investigation; **AK** - Writing – Review & Editing; **BN** – Software; **IB** – Resources; **DR** – Resources and Funding Acquisition; **DK** – Conceptualization, Project Administration, Methodology, Supervision, Writing – Review & Editing.

## Acknowledgments

We thank Maria Kurnikova for assistance in collecting samples.

We thank Ruslan Abasov for valuable comments regarding our research.

## References

1. Choi, M., Scholl, U. I., Ji, W., Liu, T., Tikhonova, I. R., Zumbo, P.,… & Nelson-Williams, C. (2009). Genetic diagnosis by whole exome capture and massively parallel DNA sequencing. Proceedings of the National Academy of Sciences, 106(45), 19096–19101.

2. Suwinski, P., Ong, C., Ling, M. H., Poh, Y. M., Khan, A. M., & Ong, H. S. (2019). Advancing personalized medicine through the application of whole exome sequencing and big data analytics. Frontiers in genetics, 10, 49.

3. Barbitoff, Y. A., Polev, D. E., Glotov, A. S., Serebryakova, E. A., Shcherbakova, I. V., Kiselev, A. M.,… & Predeus, A. V. (2020). Systematic dissection of biases in whole-exome and wholegenome sequencing reveals major determinants of coding sequence coverage. Scientific reports, 10(1), 1–13.

4. Wright, C. F., FitzPatrick, D. R., & Firth, H. V. (2018). Paediatric genomics: diagnosing rare disease in children. Nature Reviews Genetics, 19(5), 253.

5. Clark, M. J., Chen, R., Lam, H. Y., Karczewski, K. J., Chen, R., Euskirchen, G.,… & Snyder, M. (2011). Performance comparison of exome DNA sequencing technologies. Nature biotechnology, 29(10), 908–914.

6. Chilamakuri, C. S. R., Lorenz, S., Madoui, M. A., Vodák, D., Sun, J., Hovig, E.,… & Meza-Zepeda, L. A. (2014). Performance comparison of four exome capture systems for deep sequencing. BMC genomics, 15(1), 449.

7. Shigemizu, D., Momozawa, Y., Abe, T., Morizono, T., Boroevich, K. A., Takata, S.,… & Tsunoda, T. (2015). Performance comparison of four commercial human whole-exome capture platforms. Scientific reports, 5(1), 1–8.

8. García-García G. et al. Assessment of the latest NGS enrichment capture methods in clinical context //Scientific reports.--2016.--T. 6.-*N*º. 1.-C. 1–8.

9. Samorodnitsky, E., Datta, J., Jewell, B. M., Hagopian, R., Miya, J., Wing, M. R.,… & Roychowdhury, S. (2015). Comparison of custom capture for targeted next-generation DNA sequencing. The Journal of Molecular Diagnostics, 17(1), 64–75.

10. Meienberg, J., Zerjavic, K., Keller, I., Okoniewski, M., Patrignani, A., Ludin, K.,… & Matyas, G. (2015). New insights into the performance of human whole-exome capture platforms. Nucleic acids research, 43(11), e76–e76.

11. An introduction to Next-Generation Sequencing Technology https://www.illumina.com/content/dam/illumina-marketing/documents/products/illumina_sequencing_introduction.pdf

12. Fehlmann, T., Reinheimer, S., Geng, C., Su, X., Drmanac, S., Alexeev, A.,… & Keller, A. (2016). cPAS-based sequencing on the BGISEQ-500 to explore small non-coding RNAs. Clinical Epigenetics, 8(1), 1–11.

13. Chen, J., Li, X., Zhong, H., Meng, Y., & Du, H. (2019). Systematic comparison of germline variant calling pipelines cross multiple next-generation sequencers. Scientific reports, 9(1), 1–13.

14. Senabouth, A., Andersen, S., Shi, Q., Shi, L., Jiang, F., Zhang, W.,… & Powell, J. E. (2020). Comparative performance of the BGI and Illumina sequencing technology for single-cell RNA-sequencing. NAR Genomics and Bioinformatics, 2(2), lqaa034.

15. Korostin, D., Kulemin, N., Naumov, V., Belova, V., Kwon, D., & Gorbachev, A. (2020). Comparative analysis of novel MGISEQ-2000 sequencing platform vs Illumina HiSeq 2500 for whole-genome sequencing. Plos one, 15(3), e0230301.

16. Jeon, S. A., Park, J. L., Kim, J. H., Kim, J. H., Kim, Y. S., Kim, J. C., & Kim, S. Y. (2019). Comparison of the MGISEQ-2000 and Illumina HiSeq 4000 sequencing platforms for RNA sequencing. Genomics & informatics, 17(3).

17. Eberle, M. A., Fritzilas, E., Krusche, P., Källberg, M., Moore, B. L., Bekritsky, M. A.,… & Kruglyak, S. (2017). A reference data set of 5.4 million phased human variants validated by genetic inheritance from sequencing a three-generation 17-member pedigree. Genome research, 27(1), 157–164.

18. Bulusheva, I., Belova, V., Nikashin, B., & Korostin, D. (2020). BC-store: a program for mgiseq barcode sets analysis. bioRxiv.

19. MGIEasy Exome Capture V4 Probe Set User Manual https://en.mgitech.cn/Uploads/Temp/file/20191225/5e0312224c334.pdf

20. Andrews, S. (2017). FastQC: a quality control tool for high throughput sequence data. 2010.

21. Ewels, P., Magnusson, M., Lundin, S., & Käller, M. (2016). MultiQC: summarize analysis results for multiple tools and samples in a single report. Bioinformatics, 22(19), 3047–3048.

22. Bolger, A. M., Lohse, M., & Usadel, B. (2014). Trimmomatic: a flexible trimmer for Illumina sequence data. Bioinformatics, 30(15), 2114–2120.

23. Li, H., & Durbin, R. (2009). Fast and accurate short read alignment with Burrows-Wheeler transform. Bioinformatics, 25(14), 1754–1760.

24. Li, H., Handsaker, B., Wysoker, A., Fennell, T., Ruan, J., Homer, N.,… & Durbin, R. (2009). The sequence alignment/map format and SAMtools. Bioinformatics, 25(16), 2078–2079.

25. Broad Institute GitHub: http://broadinstitute.github.io/picard/

26. https://www.thermofisher.com/order/catalog/product/65001#/65001 Dynabeads™ MyOne™ Streptavidin C1: Product description

27. Wang, Q., Shashikant, C. S., Jensen, M., Altman, N. S., & Girirajan, S. (2017). Novel metrics to measure coverage in whole exome sequencing datasets reveal local and global non-uniformity. Scientific reports, 7(1), 1–11.

28. https://www.biostars.org/p/175540/

29. myBaits® Manual: https://arborbiosci.com/mybaits-manual/

30. SureSelect XT Target Enrichment for the Illumina Platform https://www.agilent.com/cs/library/usermanuals/public/G7530-90000.pdf

31. Twist Target Enrichment Protocol: https://www.twistbioscience.com/resources/protocol/twist-target-enrichment-protocol-use-twist-ngs-workflow

32. B. Faircloth, Target Enrichment of Illumina Libraries http://s3.ultraconserved.org/protocols/illumina-seqcap-hybridization-with-myselect.pdf

33. Kristina Giorda, Bahri Karaçay. Minimizing duplicates and obtaining uniform coverage in multiplexed target enrichment sequencing

34. B Marosy, J Gearhart, B Craig, KF Doheny Comparison of Whole Exome Capture Products-Coverage & Quality vs Cost. CIDR

35. Shearer, A. E., Hildebrand, M. S., Ravi, H., Joshi, S., Guiffre, A. C., Novak, B.,… & Smith, R. J. (2012). Pre-capture multiplexing improves efficiency and cost-effectiveness of targeted genomic enrichment. BMC genomics, 13(1), 1–8.

36. van der Werf, I. M., Kooy, R. F., & Vandeweyer, G. (2015). A robust protocol to increase NimbleGen SeqCap EZ multiplexing capacity to 96 samples. PloS one, 10(4), e0123872.

37. Chung, J., Son, D. S., Jeon, H. J., Kim, K. M., Park, G., Ryu, G. H.,… & Park, D. (2016). The minimal amount of starting DNA for Agilent’s hybrid capture-based targeted massively parallel sequencing. Scientific reports, 6(1), 1–10.

38. Van Dijk, E. L., Jaszczyszyn, Y., & Thermes, C. (2014). Library preparation methods for nextgeneration sequencing: tone down the bias. Experimental cell research, 322(1), 12–20.

39. Aird, D., Chen, W. S., Ross, M., Connolly, K., Meldrim, J., Russ, C.,… & Gnirke, A. (2010). Analyzing and minimizing bias in Illumina sequencing libraries. Genome biology, 11(S1), P3.

