## Supplementary material for "System Analysis of the Sequencing Quality of Human Whole Exome Samples on Bgi Ngs Platform": S1 File

### «RSMU_exome»
### Protocol of the exome enrichment of library pools for WES sequencing on MGI

#### Reagents

| | Title | Manufacturer | P/N |
|---|---|---|---|
| 1 | 80% ethanol | Constanta-Pharm M" LLC | LS-002430 |
| 2 | mQ water | Millipore | |
| 3 | Qubit dsDNA HS Assay Kit | ThermoFisher | IQ32854 |
| 4 | Qubit dsDNA BR Assay Kit | ThermoFisher | Q32853 |
| 5 | Agencourt AMPure XP | Beckman | A63881 |
| 6 | Set for Target Enrichment / SSELXT Human All Exon V7, 16 rxn | Agilent Technologies | 5191-4004 |
| 7 | Magnetic particles with streptavidin Dynabeads™ MyOne™ Streptavidin C1 (10mg/ml) | ThermoFisher | 65001 |
| 8 | COT Human DNA (500µg/packaging) | Roche | 11581074001 |
| 9 | SALMON SPERM DNA SOLN 5 X 1ML (10 mg/ml) reagent | Thermo Invitrogen | 15632011 |
| 10 | RNAase inhibitor Super Rase In, 20 units, act/mcl, 2500 units. (20U/mkl) | Thermo Invitrogen | AM2694 |
| 11 | Reagent 50X DENHARDTS SOLUTION 100ML | Thermo Invitrogen | 750018 |
| 12 | KAPA HiFi polymerase HS | KAPA biosystems | 7958897001 |
| 13 | Sodium chloride and sodium phosphate 20 X SSPE 1L | Thermo Invitrogen | AM9767 |
| 14 | Reagent 0.5 M EDTA PH 8.0 100ML (BUF KIT) | Thermo Invitrogen | AM9260G |
| 15 | 5M NACL 500 ML EACH Reagent | Thermo Invitrogen | AM9759 |
| 16 | Reagent 10% SDS SOLUTION 500 ML EACH | Thermo Invitrogen | AM9822 |
| 17 | 20X SSC BUFFER 500 ML EACH Reagent | Thermo Invitrogen | AM9770 |
| 18 | Tris-HCl 1M 7.5 pH, 1L | Thermo Invitrogen | 15567027 |
| 19 | High Sensitivity DNA reagents | Agilent Technologies | G2938-85004 |
| 20 | Agarose | | |
| 20 | Ethidium bromide | | |
| 21 | TAE buffer (x1) | Eurogen | PB022 |

| Oligonucleotides | Concentration | Sequence 5'-3 |
|---|---|---|
| MGIAd_PCR_2 | 30 µM | TGTGAGCCAAGGAGTTG |
| MGIAd_PCR_1 | 30 µM | /5Phos/GAACGACATGGCTACGA |

| | | |
|---|---|---|
| IGMblock11 | 1000 µM | GAACGACA+TGGC+TACGA+TCCGAC+TT |
| IGMblock12 | 1000 µM | TGTGAGCC+AAGG+AGTTGiiiiiiiiiiTGTCTTCCT+AAG+ACCGCTTG CCTCCG+ACTT |

#### Materials

| | Title | Manufacturer | P/N |
|---|---|---|---|
| 1 | LoBind 1.5 ml microtubes | Eppendorf | 22431021 |
| 2 | Falcon 50 ml tubes | Eppendorf | Epp 0030 122.178 |
| 3 | Filter tips 20-300 | SSI | SSI-4928NSFS |
| 4 | Filter tips 2-20 | SSI | SSI-4238NAFS |
| 5 | Handpieces with filters 1-10 | SSI | SSI-4138NSFS* |
| 6 | Filter tips 300-1200 | SSI | SSI-4348NSFS |
| 7 | Microtubes 0.5 ml | Cornig | PCR-05-C |
| 8 | Tubes 1.5ml 500 pcs/upe | SSI | SSI-1260-00 |
| 9 | SSI-3111 strips and flat caps | SSI | SSI-3105* |
| 10 | Strips Eppendorf 0.2 ml | Eppendorf | Epp 0030 124.359 |
| 11 | 0.2 PCR tubes with ind. lids | SSI | SSI |

#### Equipment

| | Title | Manufacturer | P/N |
|---|---|---|---|
| 1 | Qubit | Thermo | |
| 2 | Vortex Microspin FV-2400 | Biosan | |
| 3 | Automatic pipettes Eppendorf in the range of volumes from 0.5 to 10 µl, from 2 to 20 µl, from 200 to 200 µl, from 100 to 1000 µl | Eppendorf | |
| 4 | Racks for 1.5 ml, 0.5 ml, 0.2 ml tubes | Helicon | |
| 5 | Amplifier 96-well with manually opened lid | | |
| 6 | Orbital shaker, thermal shaker | Biosan | |
| 7 | Thermostat | DNA technology | |
| 8 | Bioanalyzer 2100 | Agilent | |
| 9 | Magnetic rack for 1.5 ml tubes and 0.2 ml tubes | Thermo, Helicon | |
| 10 | Camera for gel electrophoresis with combs, power supply, transilluminator | | |

### Exome enrichment of DNA libraries

- We use SureSelect XT1 Baits v6 or v7 (Agilent) probes. They must be stored at -70 ° C and carefully thawed on ice in the refrigerator beforehand!

- Hyb buffers #1,2,4 are stored on RT, cannot be cooled. Hyb 3 should be defrosted in advance.

- We use Blocking oligo: IGMblock11, IGMblock12. The stock concentration is 1000 µM.

#### DNA library pooling

Write down for each sample the concentration value ( ng / mkl ) measured by Qubit, select 10-13 libraries for 1 pool. In total, pool should weigh from 3 mkg to 5 mkg. The amount of each sample in the pool should be 350-500 ng (value may vary depending on the experimental goals).

***Important: Make sure that the adapters in DNA libraries do not match.***

***Important: Libraries must be made with the same DNA insert size. You can pool libraries made with size - select x0.8 + x0.2, or libraries made with size - select x0.9 + x0.2.***

It is necessary to completely dry the pool on the SpeedVac concentrator in 1.5 ml LoBind tubes at maximum vacuum and at temperature of 50°C.

#### Preparation of hybridization buffer

***Important: It* is necessary to observe the requirements for cleanliness of the workplace - before start you must wipe the work surface, pipettes and racks with 70% alcohol and leave all plastic (tips, tubes) in the box under the UV for 30 minutes.**

1. Prepare Hybridization mix for 10 pools (the volume is such for uniform heating in the thermostat and get rid of the resulting precipitate) in 1.5 ml tube:

| | x1 | x10 |
|---|---|---|
| - Hyb1 (20X SSPE) | 9 µL | 90 µL |
| - Hyb2(0.5M EDTA) | 0.5 µL | 5 µL |
| - Hyb3 (50X Denhardt's Solution) | 3.5 µL | 35 µL |
| - Hyb4 (10% SDS) | 0.5 µL | 5 µL |

Vortex well and put in a thermostat at 65 ° C to get rid of the precipitate.

#### Mixing Blocks and DNA library pools

1. Add Blocking oligos to LoBind tube with dried pool of libraries:

- Block 1 (cot DNA) = 13 µL*

**in case of reagent shortage you can reduce the volume to 10 µL*

***Important! Pipet well and peel off the walls of the mix so that the dried DNA is FULLY DISSOLVED. Vortex the tubes so that the volume of the mixture is well mixed along the lower walls of the tube.***

Transfer 13 µL pool and Block 1 mixtures to new 0.2 mL single PCR tubes. Sign ***LIBS.***
Add 500 pmol to each pool of each Blocking oligo:

- IGMblock11 = 0.5 µL
- IGMblock12 = 0.5 µL

=14 µL

2. Set the temperature program in the amplifier with a heated lid, the lid temperature should be 105 ° C. Put ***Libs*** in the thermocycler and start the program.

| Temperature | Time | Actions |
|---|---|---|
| 95°C | 05:00 | Put Libs |
| 65°C | ∞ | Put Hybs<br>Put Probes |

While the 95 ° C program is running, transfer ***15 µL of*** heated hybridization buffer into 0.2 ml tubes according to the number of pools , or it is allowed to transfer a sufficient volume for several reactions into one tube, sign ***HYBS***. Put in the thermocycler for warming up as soon as the temperature reaches 65 ° C for at least 5 minutes.

#### Probe preparation

***Important:*** Probes are of RNA nature, so the enrichment is performed with the most serious requirements to the cleanliness of the workplace, equipment and consumables.

1. Mix probes in single 0.2 mL tubes:

- Rnase Block = 1 µL
- SureSelect baits = 4 µL

2. Name the tubes ***PROBES.*** Place the tubes in a thermocycler at 65 ° C, allow to warm up for 4 minutes.

#### Hybridization of DNA probes and target areas

1. Wait until all components of the hybridization are heated up, the amplification program should switch to a constant mode of 65 ° C.
2. Open tubes directly in the thermocycler and transfer 14 µ L Hybs to 14 µ L Libs , carefully mix by pipetting.
3. Then transfer 28 µ L of Hybs + Libs to 5 µ L Probes ( sum 33 µ L ), mix gently by pipetting and close the lids.
4. ***Important! Add mineral oil to prevent samples from evaporating; oil is added to the walls of the tubes so that it gently covers the reaction mixture. Close the lids tightly.***
5. Leave hybridization for 18-24 hours;

| Temperature | The time is . | Actions |
|---|---|---|
| 65°C | ∞ | 1. Transfer 14 µL Hybs to 14 µL Libs<br><br>2. Transfer 28 µL |

| | | |
|---|---|---|
| | | Hybs+Libs to 5 µL Probes (sum = 33 µL). |

#### Day 2. Washing the enrichment reaction products

This stage should be started in advance, approximately 1.5 hours before the end of hybridization.

##### Prepare washing buffers

A. *Binding buffer*

10 mL 5 M NaCl

500 uL 1 M Tris-HCl (pH 7.5)

100 uL 0.5 M EDTA

39.4 mL ddH2O

50 mL Total Volume

B. *Wash Buffer #2*

250 uL 20X SSC

500 uL 10% SDS

49.25 mL ddH2O

50 mL Total Volume

##### Washing the enrichment reaction products

***Important!*** Place Dynabeads C1 on RT 30min before use.

***Important! All stages of washing before elution and PCR should be carried out strictly at 65°C.***

1. Prepare Wash Buffer 3 in 1.5 mL tube (per 1 pool) by mixing:

- Hyb4 = 12.1 µL

- MQ Water = 1200 µL

- Wash Buffer 2 = 300 µL

Place the tube with wash buffer into the thermostat at 65°C for 45 minutes.

2. Take 30 µL Dynabeads C1 (per 1 pool) into 1.5 mL LoBind tube, place on a magnetic stand, wait until the solution becomes transparent, remove the supernatant;
3. Remove the tube from the magnetic rack, add 200 µL Binding Buffer, Vortex+spin, place on the magnetic stand, wait 2 minutes until the solution becomes transparent, remove the supernatant;
4. Repeat step 3 twice to perform a total of 3 washes;
5. Resuspend beads in 65 µL Binding Buffer.

   ***Important:*** You can wash the Dynabeads for several samples in one tube (up to 7 samples). To do this you need to increase the buffer volume in proportion to the number of samples. For example, for 4 reactions, take 120 µL Dynabeads and wash them three times with 800 µL Binding Buffer, then resuspend to 280 µL Binding Buffer.

6. Add 5 µl/pool Block 2 (Salmon Sperm DNA 1 mg/µl) and place on an orbital shaker for 15 minutes at RT.
7. Transfer 70 µL "locked" Dynabeads to new 1.5 mL LoBind tubes, sign according to the pools.
8. Put the tubes in the thermostat at 65 ° C, warm up for 5 minutes,
9. **Open the thermocycler, transfer the entire volume of enriched pools to the Dynabeads in the thermostat, mix by pipetting;**
10. Incubate the tubes at 65°C for 30 minutes in a thermoshaker;
11. Place the tubes on a magnetic stand, wait until the solution becomes transparent (~1 min), and the magnetic particles collect at the tube wall, remove the supernatant;
12. Add 500 µL of preheated Wash Buffer 3, Vortex+spin, put in a thermoshaker at 65°C for 10 minutes.
13. Carefully centrifuge the drops, transfer tubes to a magnetic stand, remove the supernatant;
14. Repeat steps 12-13 two more times (total 3 washes with WB3);
15. Dry beads on a magnetic stand for 3-4 minutes.
16. Resuspend in 31 µL MQ.

#### Post-capture PCR

At this stage, enriched libraries are "detached" from probes during denaturation and then amplified.

**IMPORTANT: Before setting up the PCR, you must first "remove" = denaturate DNA from Dynabeads.** To do this, transfer the entire reaction volume (31 µl) into the new 0.2 µl PCR tubes, put in the thermocycler at 95 ° C for 5 minutes. Then quickly put on a magnetic stand and transfer the transparent supernatant into the new 1.5 mL LoBind tube.

In the PCR use half of the enriched sample (15-17 µl), sign the 2nd half and store at -20 ° C.

Use reagents from KAPA HiFi HotStart PCR Kit.

**Important: For PCR after enrichment the common reagent should NOT be used. Buffer, dNTPs and primer aliquots must be proprietary for WES only.**

*Primers used (at a concentration of 30 µM):*

*MGIAd_PCR_1* *5Phos/GAACGACATGGCTACGA*
*MGIAd_PCR_2* *TGTGAGCCAAGGAGTTG*

1. Mix the following components in a sterile PCR tube:

- KAPA HiFi GC Buffer (5X) = 10 µL
- KAPA dNTP Mix = 1.5 µL

- MGI primer 1 = 8 µL
- MGI primer 2 = 8 µL
- KAPA HiFi HotStart DNA Polymerase = 1 µL
- DNA enriched fragments = 15-17 µL
- mQ = up to 50 µL

Total: 50 µL

1. Put into the thermocycler and start the following program:

| Temperature | The time is . | Repeats |
|---|---|---|
| 95°C | 03:00 | |
| 98°C | 00:20 | X10* |
| 60°C | 00:15 | |
| 72°C | 00:30 | |
| 72°C | 10:00 | |
| 4°C | Storage | |

*NB! The number of PCR cycles may vary depending on the use of probes from different manufacturers. For 4µl Agilent, 10 cycles are optimal.

**2.** Select 2.5 µl of PCR product and immediately perform Gel-Electrophoresis (2% agarose gel, 30 min 130 V) for quality control. A good result is a length distribution with a maximum in the range of 300-500 bp.

***Important! Step 3 is critical, if the gel is empty, it makes sense to reanimate the situation by adding a PCR mix and run few more PCR cycles.***

**3.** If the gel is all OK, then proceed to further purification of the PCR product.

***Stopping point***: if processed the next day samples can be storeat +4°C overnight.

#### Purification using Agencourt AMPure XP)

1. Transfer the reaction to a new 1.5 ml LoBind tube
2. Add to 50 µL reaction 50 µL Ampure beads (**1x** of reaction volume), vortex+spin;
3. Incubate at room temperature for 7 minutes;
4. Transfe the tubes to a magnetic rack. After the mixture becomes transparent (it takes 7 minutes), you should carefully remove the supernatant without touching the beads that contain fragments of DNA;
5. Add 400 µL fresh (!) 80% EtOH, rotate the tube in a rack 180° so that the beads run through the EtOH solution several times, collect all the beads on one wall, gently remove EtOH without touching the bids;
6. Repeat step 5 one more time to get 2 washes;
7. Remove residues of 80% EtOH with a pipette with 2-20 tips. Leave the test tubes with an open lid to dry directly on the magnetic stand for 3-5 minutes, making sure that the beads do not dry out (when dried they lighten).
8. Add 38 µL mQ, resuspend beads well on a Vortex+Spin, incubate for 7 min, then place on a magnetic stand, incubate for 7 min. Transfer the supernatant to a new tube. The supernatant contains purified DNA.

#### Enrichment quality control

To carry out the quantitative control of the obtained enriched pools - to measure the concentration with the Qubit High Sense set according to the manufacturer's protocol.

The total amount of enriched pool should be **at least 70 ng. If the amount is less, it is necessary to pool with other samples in the further reaction of circularization to obtain sufficient amount of DNA for circulation.**

Optionally – perform quality controlof the obtained enriched pools on the Agilent Bioanalyzer 2100 using the High Sensitivity Kit according to the manufacturer's protocol.

A good result is a length distribution with a peak of 250-400 bp. A good pool suitable for further preparation for sequencing is shown in *Figure 1*.

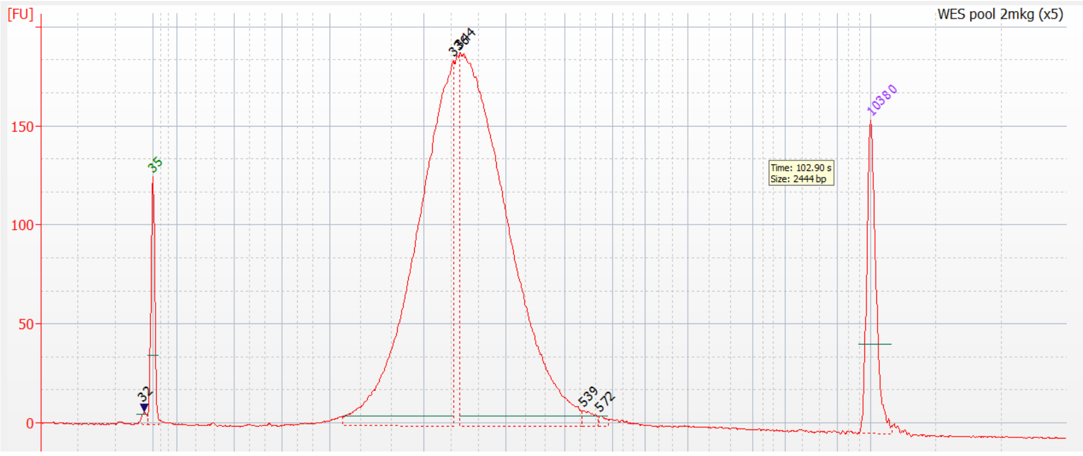


*Fig.1. Chromatogram of enriched DNA library with 350 bp peak for MGI sequencing.*
