## Supplementary material for "System Analysis of the Sequencing Quality of Human Whole Exome Samples on Bgi Ngs Platform": S2 File: marked_AUV193_1B_OL_v6_fastqc.html

marked\_AUV193\_1B\_OL\_v6.bam FastQC Report 

FastQC Report

Mon 12 Oct 2020  
marked\_AUV193\_1B\_OL\_v6.bam

### Summary

- Basic Statistics
- Per base sequence quality
- Per sequence quality scores
- Per base sequence content
- Per sequence GC content
- Per base N content
- Sequence Length Distribution
- Sequence Duplication Levels
- Overrepresented sequences
- Adapter Content

### Basic Statistics

| Measure | Value |
| --- | --- |
| Filename | marked\_AUV193\_1B\_OL\_v6.bam |
| File type | Conventional base calls |
| Encoding | Sanger / Illumina 1.9 |
| Total Sequences | 75698827 |
| Sequences flagged as poor quality | 0 |
| Sequence length | 30-150 |
| %GC | 51 |

### Per base sequence quality

### Per sequence quality scores

### Per base sequence content

### Per sequence GC content

### Per base N content

### Sequence Length Distribution

### Sequence Duplication Levels

### Overrepresented sequences

No overrepresented sequences

### Adapter Content

Produced by FastQC (version 0.11.9)
