## Supplementary figures and images for "System Analysis of the Sequencing Quality of Human Whole Exome Samples on Bgi Ngs Platform"

### S1 Figure

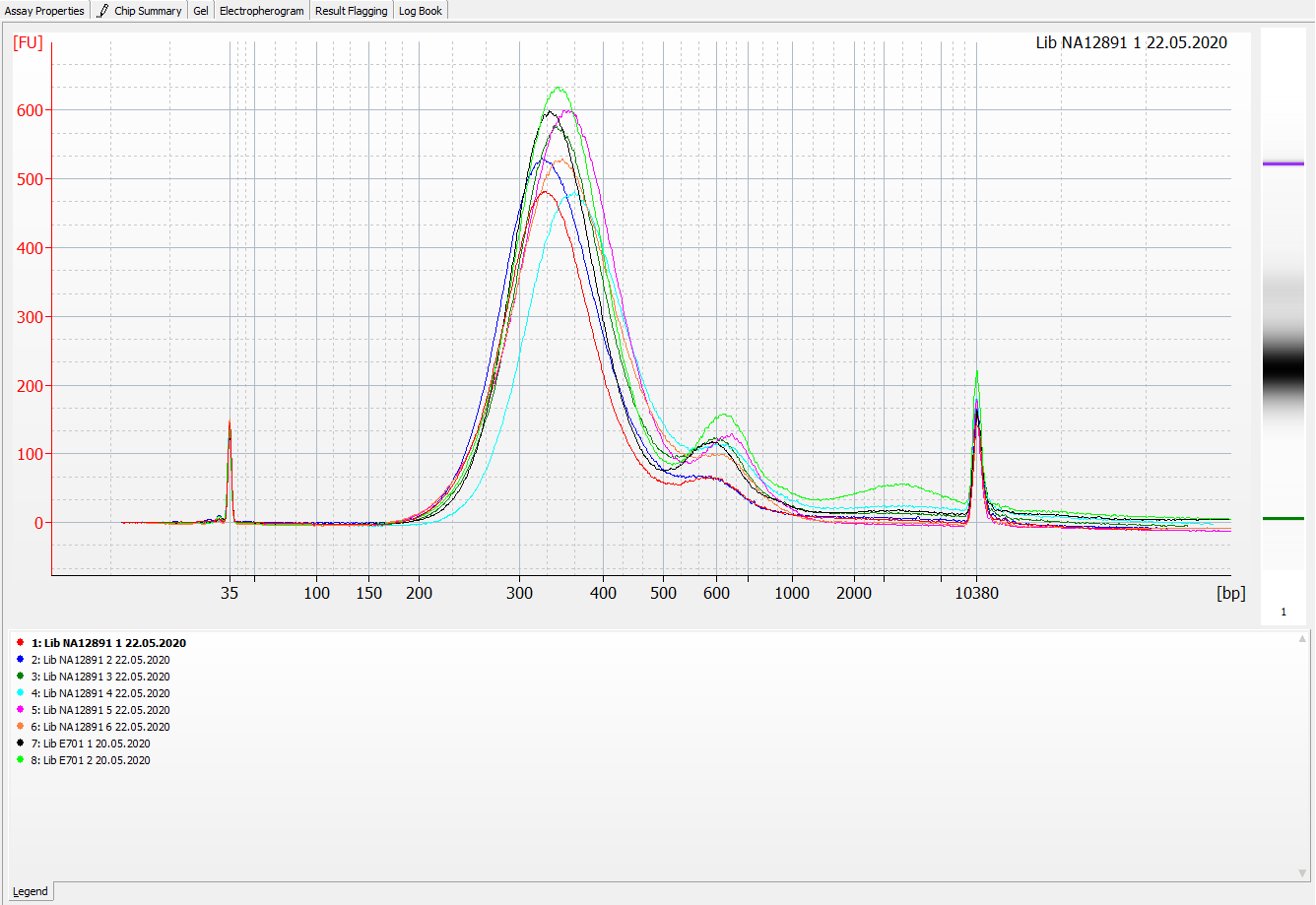
